# PRC1 directs PRC2-H3K27me3 deposition to shield adult spermatogonial stem cells from differentiation

**DOI:** 10.1101/2023.11.16.567444

**Authors:** Mengwen Hu, Yu-Han Yeh, So Maezawa, Toshinori Nakagawa, Shosei Yoshida, Satoshi H. Namekawa

## Abstract

Spermatogonial stem cell functionality resides in the slow-cycling and heterogeneous undifferentiated spermatogonia cell population. This pool of cells supports lifelong fertility in adult males by balancing self-renewal and differentiation to produce haploid gametes. However, the molecular mechanisms underpinning long-term stemness of undifferentiated spermatogonia during adulthood remain unclear. Here, we discover that an epigenetic regulator, Polycomb repressive complex 1 (PRC1), shields adult undifferentiated spermatogonia from differentiation, maintains slow cycling, and directs commitment to differentiation during steady-state spermatogenesis in adults. We show that PRC2-mediated H3K27me3 is an epigenetic hallmark of adult undifferentiated spermatogonia. Indeed, spermatogonial differentiation is accompanied by a global loss of H3K27me3. Disruption of PRC1 impairs global H3K27me3 deposition, leading to precocious spermatogonial differentiation. Therefore, PRC1 directs PRC2-H3K27me3 deposition to maintain the self-renewing state of undifferentiated spermatogonia. Importantly, in contrast to its role in other tissue stem cells, PRC1 negatively regulates the cell cycle to maintain slow cycling of undifferentiated spermatogonia. Our findings have implications for how epigenetic regulators can be tuned to regulate the stem cell potential, cell cycle, and differentiation to ensure lifelong fertility in adult males.

## Introduction

Mammalian males sustain lifelong fertility relying on balanced self-renewal and differentiation of spermatogonial stem cells (SSCs), which are destined to produce a massive number of haploid spermatozoa; ∼1,000 sperm per second for decades in average human males (1). In adult mice, stem cell activity resides in a heterogenous population of slow-cycling undifferentiated type A spermatogonia (A_undiff_, including A_single_, A_paired,_ and A_aligned_ spermatogonia), whose cell states are maintained at a delicate equilibrium (2,3). A fraction of A_undiff_ that are positive for the cell surface receptor GFRα1 comprises the vast self-renewing compartment in homeostatic, steady-state spermatogenesis. While A_undiff_ expressing NGN3 (Neurog3), which are GFRα1-negative, are destined for differentiation but retain the capacity to revert to the GFRα1^+^ state and repopulate the self-renewing pool (Figure 1A) (4,5). In response to retinoic acid (RA) signaling, the NGN3^+^ fraction of A_undiff_ undergo irreversible commitment to become fast-cycling KIT^+^ differentiating spermatogonia (designated A_1_, A_2_, A_3_, A_4_, Intermediate (In), and B spermatogonia), which are devoid of stem cell potential (Figure 1A) (6) and undergo a series of mitotic divisions before entering meiosis (7,8). Therefore, the transition from A_undiff_ (KIT^−^) to KIT^+^ differentiating spermatogonia is accompanied by the irreversible loss of self-renewing stem cell potential and the acceleration of their cell cycle.

**Figure 1.**
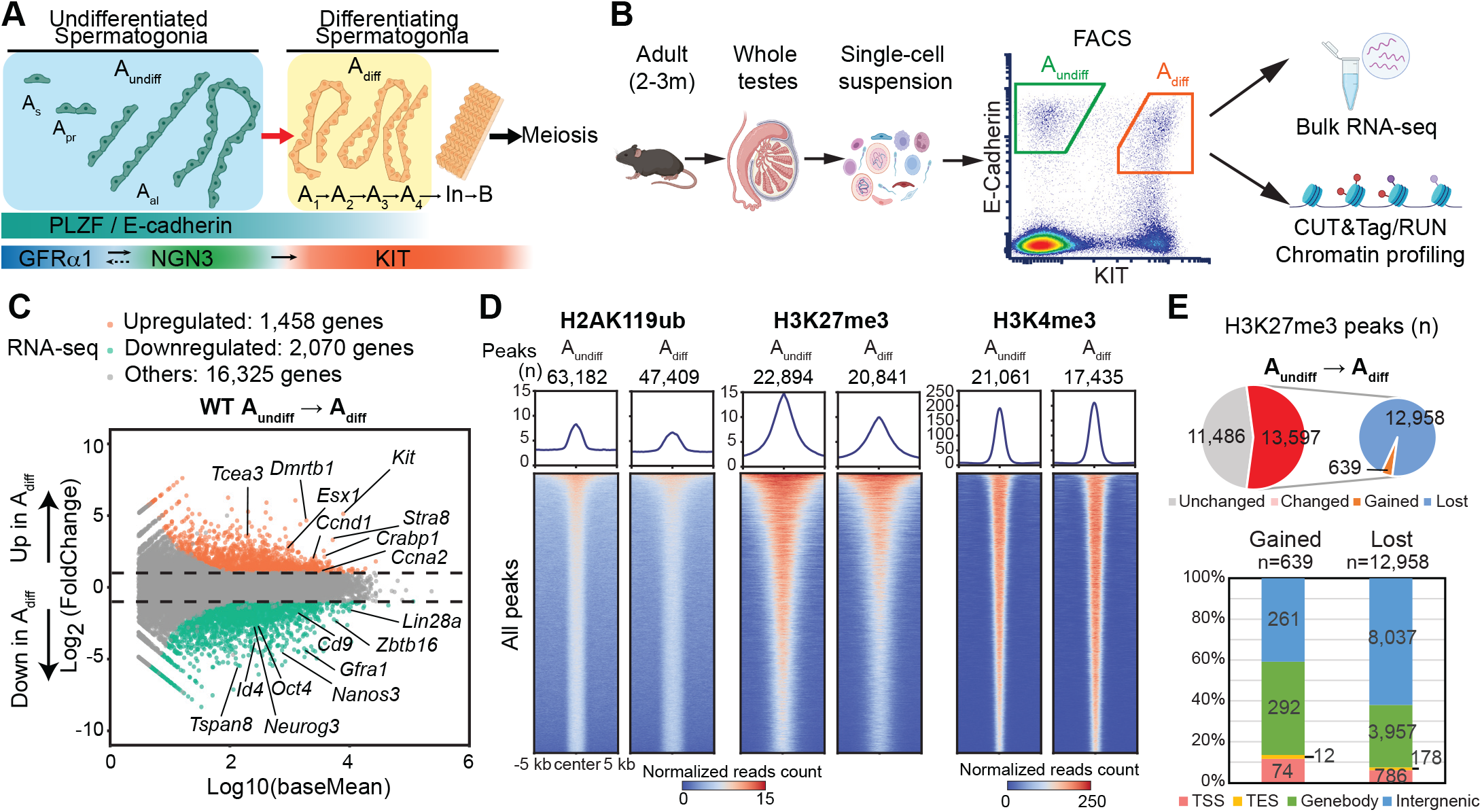
Dynamic transcriptomic and epigenomic changes during spermatogonial differentiation in adult wild-type mice. (A) Schematic of spermatogonial hierarchy and marker gene expression. Based on nuclear morphology, spermatogonia can be classified as type A, intermediate, or type B. Type A includes the undifferentiated spermatogonia (A_undiff_, including A_single_ (A_s_), A_paired_(A_pr_), and A_aligned_(A_al_)) and early differentiating spermatogonia (A_diff_, including A_1_, A_2_, A_3,_ and A_4_), whereas intermediate (In) and type B encompass the later differentiating spermatogonia. (B), Overview of the experimental steps to perform bulk RNA-Seq and chromatin profiling using CUT&RUN/Tag on FACS-isolated spermatogonia. (C) MA plot showing a comparison of transcriptomes between A_undiff_ and A_diff_ in adult WT mice (2-3 mo). All genes with False Discovery Rate (FDR) values are plotted. Differentially expressed genes (DEGs: Log_2_FoldChange >1, FDR< 0.05) are color-coded (orange: upregulated genes; green: downregulated genes). Two independent biological replicates were examined for RNA-seq. (D) Heatmaps and average tag density plots showing peaks of H2AK119ub, H3K27me3, and H3K4me3 in WT A_undiff_ and A_diff_. The numbers of peaks are indicated (n). The color keys represent signal intensity, and the numbers represent spike-in normalized reads counts. (E) Pie chart and bar chart showing the H3K27me3 peak change and the genomic distribution of gained and lost H3K27me3 peaks during spermatogonial differentiation. The numbers of peaks are shown (n).

Spermatogenesis thus provides a unique *in vivo* model for studying robust and persistent tissue stem cell systems. Previous studies have identified the regulatory mechanisms, including key signaling pathways and transcription factors, that support the formation and maintenance of undifferentiated spermatogonia (9–11). For example, growth factors produced within the testicular microenvironment, such as GDNF and FGFs, promote SSC self-renewal (12–14). GDNF acts through the cell surface GFRα1/RET receptor complex and activates the downstream PI3K/AKT, RAS/ERK MAPK and the Src family kinase (SFK) pathways, leading to the maintenance of the undifferentiated state of SSCs (15–17). In contrast, RA induces spermatogonial differentiation; depletion of RA results in the accumulation of A_undiff_ (18,19). In addition to these extrinsic stimuli, various cell-intrinsic factors are required to maintain undifferentiated spermatogonia, including the transcription factors PLZF (20–22), SALL4 (23,24), ETV5 (25), DMRT1 (26), and FOXC2 (27); RNA-binding proteins NANOS2 (28) and DND1 (29); epigenetic regulators Polycomb repressive complexes (30–32) and DOT1L (33); DNA damage response proteins ATM (34) and FANCB (35); the tumor suppressor RB (36), and SHISA6, an inhibitor of differentiation-promoting Wnt/β-Catenin signaling (8).

However, while numerous factors have been linked to the formation and maintenance of undifferentiated spermatogonia, major questions remain regarding the molecular underpinnings that shield and maintain robust stemness in these cells. In particular, it is unclear how slow cycling is regulated while lifelong differentiation potential is maintained in adult undifferentiated spermatogonia. This is primarily due to the fact that most previous studies have focused on juvenile spermatogenesis before the establishment of steady-state adult spermatogenesis (37,38) or used cultured germline stem cells that do not fully represent *in vivo* features of spermatogonial stem cells (39,40).

Epigenetic mechanisms that generate long-lasting chromatin states and define gene expression programs are critical for the development and regulation of stem cells (41). Polycomb group proteins (PcGs) mediate such mechanisms by establishing repressive chromatin and defining cell-type specific gene expression programs (42,43). Two major functionally-related Polycomb repressive complexes— PRC1 and PRC2— catalyze the formation of two major repressive histone marks, monoubiquitination of H2A at lysine 119 (H2AK119ub) and trimethylation of H3 at lysine 27 (H3K27me3) (44), respectively. Analysis of PcG-mediated modifications has provided a picture of the epigenomic landscape of male germ cells, including undifferentiated spermatogonia (37,38,45–47). Of note, undifferentiated spermatogonia contain bivalent chromatin; the repressive H3K27me3 modification co-occurs with an active mark, H3 trimethylated at lysine 4 (H3K4me3) (37,45), a feature associated with developmental potential in the germline. Nevertheless, the functions of these marks remain largely speculative due to the lack of quantitative analysis combined with genetic approaches.

In this study, by quantitative epigenomic analysis, we define H3K27me3 as a major barrier to spermatogonial differentiation, arguing that this is a hallmark of the undifferentiated state in adult undifferentiated spermatogonia. Indeed, spermatogonial differentiation is accompanied by the global loss of H3K27me3. By using a PRC1 conditional loss-of-function model, we show that PRC1 directs deposition of PRC2-mediated H3K27me3 to shield the undifferentiated state and control spermatogonial differentiation. Unlike other tissue stem cell systems where PcG proteins promote cell cycle progression, we demonstrate that PRC1 acts as a negative cell cycle regulator to ensure the slow cycling property of undifferentiated spermatogonia. Therefore, our study uncovers that key features of undifferentiated spermatogonia, including the undifferentiated state, slow cycling property, and the potential for commitment to differentiation, are molecularly linked by PRC1.

## Materials and Methods

### Animals

Generation of conditionally deficient *Rnf2* mice in a *Ring1^-/-^* background was performed as described previously (30). Briefly, PRC1cKO mice *Ring1^-/-^; Rnf2^F/-^; Ddx4-Cre* were generated from *Ring1^-/-^; Rnf2^F/F^* females crossed with *Ring1^-/-^; Rnf2^F/+^; Ddx4-Cre* males and PRC1ctrl mice used in experiments were *Ring1^-/-^; Rnf2^F/+^; Ddx4-Cre* littermate males. Mice were maintained on a mixed genetic background of FVB and C57BL/6J. Generation of mutant *Ring1* (*Ring1^-/-^*) and *Rnf2* floxed alleles (*Rnf2^F/F^*) was reported previously(48,49). *Ring1^-/-^* and *Rnf2^F/F^*mice were obtained from Dr. Haruhiko Koseki. *Ddx4-Cre* transgenic mice and wild-type (WT) C57BL/6J mice were purchased from the Jackson Laboratory (50). For each experiment, a minimum of three independent mice were analyzed. Mice were maintained on a 12:12 hour light: dark cycle in a temperature and humidity-controlled vivarium (22 ± 2 °C; 40–50% humidity) with free access to food and water in the pathogen-free animal care facility. Mice were used according to the guidelines of the Institutional Animal Care and Use Committee (IACUC: protocol no. IACUC2018-0040 and 21931) at Cincinnati Children’s Hospital Medical Center and the University of California, Davis.

### Win18,446 treatment

For Win18,446 injection experiments, 1-month-old male mice were injected subcutaneously with 40 μg/μl Win18,446 (N, N’-Octamethylenebis(2,2-dichloroacetamide), MP Biomedicals, 02158050), dissolved in 50 μl dimethyl sulfoxide (DMSO). After ∼30 consecutive daily treatments, mice were euthanized 24h post-treatment for sampling. Collected testes were either fixed for histological analyses or dissociated for cell sorting.

### Histology and Immunostaining

For the preparation of testicular paraffin blocks, testes were fixed with 4% paraformaldehyde (PFA) overnight at 4°C, followed by dehydration and embedding in paraffin. For immunostaining, 7-µm-thick paraffin sections were deparaffinized and autoclaved in target retrieval solution (DAKO) for 10 min at 121°C. Sections were blocked with Blocking One Histo (Nacalai) for 1 h at room temperature and then incubated with primary antibodies as outlined below: goat anti-Human PLZF (1:200, R&D, AF2944), Rabbit anti-Ki67 (1:100, Abcam, ab16667). Sections were washed with PBST (PBS containing 0.1% Tween 20) three times at room temperature for 5 min and then incubated with the corresponding secondary antibodies (Donkey Anti-Goat IgG (H+L) Alexa Fluor 488, A-11055; Donkey Anti-Rabbit IgG (H+L) Alexa Fluor 555, A-31572; Invitrogen) at 1:500 dilution for 1h at room temperature. Finally, sections were counterstained with DAPI and mounted using 20 µL undiluted ProLong Gold Antifade Mountant (ThermoFisher Scientific, P36930).

Similarly, sorted cells were fixed with 4% PFA and permeabilized with 0.5% Triton-X in PBS, followed by blocking with 1% BSA and 4% Donkey Serum (Sigma, D9663) in PBST for 1 h at room temperature. Cells were then incubated with the following primary antibodies: goat anti-Human PLZF (1:200, R&D, AF2944), or goat anti-CD117/c-kit (1:200, R&D, AF1356), and rabbit anti-DDX4 (1:400, Abcam, ab13840). After overnight incubation at 4°C, cells were washed and incubated with secondary antibodies (Donkey Anti-Goat IgG (H+L) Alexa Fluor 488, A-11055; Donkey Anti-Rabbit IgG (H+L) Alexa Fluor 555, A-31572; Invitrogen) at 1:500 dilution for 1h at room temperature. Finally, cells were counterstained with DAPI and mounted onto slides using ProLong Gold Antifade Mountant. Images were obtained by confocal laser scanning microscopy (A1R, Nikon) and processed with NIS-Elements (Nikon).

### Flow cytometry and cell sorting

Flow cytometric experiments and cell sorting were performed using SH800S (SONY), with antibody-stained testicular single-cell suspensions prepared as described previously (51). Data were analyzed with SH800S software (SONY) and FCS Express 7 (De Novo Software).

In brief, to prepare single cells suspension for cell sorting, detangled seminiferous tubules from adult mouse testes were first washed with 1x Hanks’ Balanced Salt Solution (HBSS, Gibco, 14175095) three times and then incubated in Dulbecco’s Modified Eagle Medium (DMEM, Gibco, 11885076) containing 2% Fetal Bovine Serum (FBS, Gibco, 16000044), 2 mg/ml Type 1 collagenase (Worthington, CLS1), 0.25 mg/ml DNase I (Sigma, D5025), 1.5 mg/ml Hyaluronidase (Sigma, H3506) and 700 U/ml Recombinant Collagenase (FUJIFILM Wako Pure Chemical Corporation, 036-23141) at 37℃ for 20 min, then dissociated using vigorous pipetting and incubated at 37℃ for another 10 min. The single cell suspension was then washed with 10 mL FACS buffer (PBS containing 2% FBS) three times by centrifugation at 300 × g for 5 minutes and filtered through a 70 µm nylon cell strainer (Falcon, 352350).

The resultant single cells were stained with cocktails of antibodies diluted with FACS buffer listed as follows: PE-conjugated anti-mouse/human CD324 (E-Cadherin) antibody (1:500, Biolegend, 147303), PE/Cy7-conjugated anti-mouse CD117 (c-Kit) antibody (1:200, Biolegend, 105814) and FITC-conjugated anti-mouse CD9 antibody (1:500, Biolegend, 124808). After 1h incubation on ice, cells were washed with 10 mL FACS buffer three times by centrifugation at 300 × g for 5 minutes and filtered into a 5 mL FACS tube through a 35 µm nylon mesh cap (Falcon, 352235). 7-AAD Viability Stain (Invitrogen, 00-6993-50) was added to cell suspension for the exclusion of dead cells. Samples were kept on ice until sorting.

Cells were analyzed after removing small and large debris in FSC-A versus SSC-A gating, doublets in FSC-W versus FSC-H gating, and 7AAD+ dead cells. Then the desired cell population was collected in gates determined based on antibody staining (Supplementary Figure S1A).

### RNA-seq library generation and sequencing

RNA-seq libraries from PRC1ctrl and PRC1cKO A_undiff_ were prepared as follows using the Smart-Seq2 method. ∼7,000 A_undiff_ cells were isolated from 2-month-old mice and pooled as one replicate. Two independent biological replicates were used for RNA-seq library generation. Total RNA was extracted using the RNeasy Plus Micro Kit (QIAGEN, Cat # 74034) according to the manufacturer’s instructions. Libraries were constructed using NEBNext® Single Cell/Low Input RNA Library Prep Kit for Illumina® (NEB, E6420S) according to the manufacturer’s instructions. Prepared RNA-seq libraries were sequenced on the Novaseq 6000 system (Illumina) with 100-bp paired-end reads.

Stranded RNA-seq libraries were prepared for (1) WT A_undiff_ and WT A_diff_; (2) PRC1ctrl A_undiff_ and PRC1cKO A_undiff_ after Win18,446 treatment. ∼100,000-150,000 cells isolated from adult mice were pooled as one replicate, and two independent biological replicates were used for RNA-seq library generation. Total RNA was extracted using the RNeasy Plus Mini Kit (QIAGEN, Cat # 74134) according to the manufacturer’s instructions. Library construction was performed by the CCHMC DNA Sequencing and Genotyping Core (Cincinnati, Ohio, USA) using the Illumina TruSeq stranded mRNA kit after polyA enrichment. Prepared RNA-seq libraries were sequenced on the Novaseq 6000 system (Illumina) with 100-bp paired-end reads.

### CUT&RUN/Tag library generation and sequencing

CUT&RUN libraries of PRC1ctrl, PRC1cKO, and WT spermatogonia were conducted as previously described (52) (a step-by-step protocol https://www.protocols.io/view/cut-amp-run-targeted-in-situ-genome-wide-profiling-14egnr4ql5dy/v3). ∼10,000 FACS-sorted cells were pooled as one replicate, and two independent biological replicates were used for CUT&RUN library generation. The antibodies used were rabbit anti-H3K4me3 (1/100; Cell Signaling Technology; 9751), and rabbit anti-H3K27me3 antibody (1/100; Cell Signaling Technology; 9733). Homemade Protein A/G-MNase was provided by Dr. Artem Barski’s laboratory. Libraries were constructed using NEBNext® Ultra II DNA Library Prep Kit for Illumina (NEB, E7645S). Prepared CUT&RUN libraries of PRC1ctrl and PRC1cKO A_undiff_ were sequenced on the NovaSeq 6000 system (Illumina) with paired-ended 50-bp reads, and WT CUT&RUN libraries for H3K4me3 were sequenced on the NextSeq 500 system (Illumina) with paired-ended 75-bp reads.

CUT&Tag libraries of WT A_undiff_ and A_diff_ for H2AK119ub and H3K27me3 were prepared as previously described(53,54) (a step-by-step protocol https://www.protocols.io/view/bench-top-cut-amp-tag-kqdg34qdpl25/v3) using CUTANA™ pAG-Tn5 (Epicypher, 15-1017). Quantitative spike-in CUT&Tag was performed by adding Drosophila S2 cells at a 2:1 ratio to mouse spermatogonial cells (20,000 S2 cells to 10,000 mouse cells) in each reaction. The antibodies used were rabbit anti-H2AK119ub (1/100; Cell Signaling Technology; 8240) and rabbit anti-H3K27me3 antibody (1/50; Cell Signaling Technology; 9733). CUT&Tag libraries were sequenced on the HiSeq 4000 system (Illumina) with 150-bp paired-end reads.

### RNA-seq data processing

Raw RNA-seq reads after trimming by Trim-galore (https://github.com/FelixKrueger/TrimGalore) (version 0.6.6) with the parameter ‘--paired’ were aligned to the mouse (GRCm38/mm10) genome using HISAT2 (55) (version 2.2.1) with default parameters. All unmapped reads, non-uniquely mapped reads, and reads with low mapping quality (MAPQ < 30) were filtered out by samtools (56) (version 1.9) with the parameter ’-q 30’ before being subjected to downstream analyses.

To identify differentially expressed genes in RNA-seq, raw read counts for each gene were generated using the htseq-count function, part of the HTSeq package (57) (version 1.6.0) based on mouse gene annotations (gencode.vM24.annotation.gtf, GRCm38/mm10). DESeq2 (58) (version 1.28.1) was used for differential gene expression analyses with cutoffs log2FoldChange > 1 and FDR values (P adjusted: P_adj_ values) < 0.05. In DESeq2, P values attained by the Wald test were corrected for multiple testing using the Benjamini and Hochberg method by default (FDR values). FDR values were used to determine significantly dysregulated genes. The TPM values of each gene were generated using RSEM (59) (version 1.3.3) for comparative expression analyses and computing the Pearson correlation coefficient between biological replicates.

To perform GO analyses, we used the online functional annotation clustering tool Metascape (60) (http://metascape.org). Bioinformatics analyses were visualized as heatmaps using Morpheus (https://software.broadinstitute.org/morpheus/). The TPM values of each gene were first transformed by the Robust Z-Score Method, also known as the Median Absolute Deviation method, or log-transformed, then used as input for plotting. PCA plot was generated using TPM values by the web tool ClustVis (https://biit.cs.ut.ee/clustvis/).

### CUT&RUN/Tag data processing

For CUT&RUN/Tag data processing, we basically followed the online tutorial posted by the Henikoff Lab (https://protocols.io/view/cut-amp-tag-data-processing-and-analysis-tutorial-bjk2kkye.html). Briefly, after trimming by Trim-galore (https://github.com/FelixKrueger/TrimGalore) (version 0.6.6), raw paired-end reads were aligned with the mouse genome (GRCm38/mm10) using Bowtie2 (61) (version 2.4.2) with options: --end-to-end --very-sensitive --no-mixed --no-discordant -- phred33 -I 10 -X 700. Bacterially-produced pAG-MNase protein brings along a fixed amount of *E. coli* DNA that can be used as spike-in DNA for CUT&RUN. *D. melanogaster* DNA delivered by *Drosophila* S2 cells was used as spike-in DNA for CUT&Tag. For mapping *E.coli* or *D. melanogaster* spike-in fragments, we also use the --no-overlap --no-dovetail options to avoid cross-mapping. PCR duplicates were removed using the ‘MarkDuplicates’ command in Picard tools (version 2.23.8) (https://broadinstitute.github.io/picard/). To compare replicates, Pearson correlation coefficients were calculated and plotted by multiBamSummary bins and plot correlation from deepTools (62) (version 3.5.0). Biological replicates were pooled for visualization and other analyses after validation of reproducibility (Pearson correlation coefficient > 0.85). Spike-in normalization was implemented using the exogenous scaling factor computed from the *E.coli* or dm6 mapping files (scale factors = 100,000/ spike-in reads for CUT&RUN or 1,000,000 / spike-in reads for CUT&Tag).

SEACR (63) (https://seacr.fredhutch.org/) was used for peak calling, and HOMER (64) (version 4.11) was used for peak annotation. For visualization using the Integrative Genomics Viewer (65) (version 2.5.3), spike-in normalized genome coverage tracks were generated using the genomecov function of bedtools (66) (version 2.29.2) with the ‘-bg -scale $scale_factor’ parameter. K-means clustering analysis was performed using seqMINER (67) (version 1.3.4). DeepTools (62) was used to draw tag density plots and heatmaps for read enrichments. Differential enrichment analysis was conducted using the R package DiffBind (68) (version 3.6.5) with spike-in normalization mode. The FPKM values of enrichment for each gene’s promoter region (TSS ± 2kb) were generated using Bamscale package (69) (version 0.0.5) and normalized by scaling factor for comparative enrichment analyses.

Gene Ontology Biological Process (GO-BP) term enrichment and ChIP-x Enrichment Analysis (ChEA) using the gene sets clustered from the CUT&RUN/Tag results was performed using the Enricher website (https://maayanlab.cloud/Enrichr/) (70,71). A dot plot was produced using the R package ggplot2 (3.3.6).

### CUT&RUN and ChIP-seq data reanalysis

Processed bigwig files of CBX2, PHC2, and BMI1 CUT&RUN in adult spermatogonia and bedGraph files of RAR ChIP-seq in GS cells were downloaded from Gene Expression Omnibus (GEO) under accession no. GSE210367 and GSE116798, respectively, and used for plotting directly. Raw STRA8 ChIP-seq data of mouse testes were downloaded from the DDBJ Sequence Read Archive (DRA) under accession number DRA007778 and aligned with the mouse genome (GRCm38/mm10) using Bowtie2 with default options. Aligned bam files were converted to bigwig files using Deeptools by normalizing with RPKM and then used for plotting.

### Statistics

Statistical methods and P values for each plot are listed in the figure legends and/or in the Methods. Statistical significance for pairwise comparisons was determined using two-tailed unpaired t-tests. Next-generation sequencing data (RNA-seq, CUT&RUN/Tag) were based on two independent replicates. No statistical methods were used to predetermine sample size in these experiments. Experiments were not randomized, and investigators were not blinded to allocation during experiments and outcome assessments.

## Results

### Dynamic gene expression changes during spermatogonial differentiation

To elucidate how A_undiff_ are maintained and how they lose stem cell potential and commit to differentiation during adult spermatogenesis, we first isolated A_undiff_ and early differentiating spermatogonia and performed gene expression analysis using bulk RNA-seq. We isolated these fractions from WT adult male C57BL/6 mice at 2-3 months (mo) of age using our previously established fluorescence-activated cell sorting (FACS) method (Figure 1B, Supplementary Figure S1A) (2). We utilized the cell surface protein E-Cadherin to isolate A_undiff_ and early differentiating spermatogonia (72,73), and KIT expression to distinguish differentiating spermatogonia (Figure 1A). In addition, another cell surface protein, CD9, enabled the isolation of a highly pure spermatogonial population while excluding somatic contaminants (73). Using this method, A_undiff_ were collected based on E-Cadherin expression and high levels of CD9, as well as the absence of KIT (E-Cadherin^+^CD9^high^KIT^−^); early differentiating spermatogonia were isolated based on the expression of KIT and E-Cadherin, and a medium level of CD9 (E-Cadherin^+^CD9^medium^KIT^+^). The E-Cadherin^+^CD9^medium^KIT^+^ fraction contains highly enriched early stages of differentiating spermatogonia, most likely A_1_ through A_4_ spermatogonia, rather than In and B spermatogonia that are about to enter meiosis (2,72,74) (Supplementary Figure S1A). Therefore, we hereafter abbreviate the E-Cadherin^+^CD9^medium^KIT^+^ fraction as A_diff_. We confirmed the high purity of these cell populations (Supplementary Figure S1B-C) and high reproducibility of RNA-seq data between biological replicates (Supplementary Figure S1D).

Comparing gene expression between A_undiff_ and A_diff_ revealed a dynamic change in the transcriptome during spermatogonial differentiation, marked by the upregulation of 1,458 genes and downregulation of 2,070 genes (Figure 1C, Supplementary Data 1). GO term analysis revealed that genes involved in cell morphogenesis, cell cycle, cell fate commitment, and meiosis were upregulated upon differentiation. In contrast, genes functioning in cytokine production, cell adhesion, and especially stem cell population maintenance were downregulated (Supplementary Figure S1E). Consistent with this analysis, A_undiff_ markers (*Id4*, *Gfra1*, *Zbtb16*(*PLZF*), *Neurog3*(*Ngn3*), and *Tspan8*) and pluripotency markers (*Oct4*, *Nanos3,* and *Lin28a*) were downregulated upon differentiation, while *Kit* was upregulated in A_diff_ (Figure 1C). In addition, *Stra8* and retinoic-acid-binding-protein 1 *Crabp1* were significantly upregulated in A_diff_ (Figure 1C), consistent with the roles of RA signaling in spermatogonial differentiation. Further, cell cycle factors (*Ccnd1*, *Ccna2*, *Ccne1,* and *Ccne2*) were markedly upregulated in A_diff_, in line with the rapid proliferative state of this population, and other recently-identified markers for spermatogonial differentiation (such as *Esx1* and *Tcea3*)(75) were also upregulated in A_diff_ (Figure 1C, Supplementary Figure S1F). Taken together, these analyses confirm that distinct gene expression programs are active in A_undiff_ compared to A_diff_ populations in adult testes.

### Polycomb-mediated chromatin remodeling underlies spermatogonial differentiation

To determine the chromatin states underlying the gene expression changes during spermatogonial differentiation, we performed CUT&Tag (Cleavage Under Targets and Tagmentation)(53,54) or CUT&RUN (cleavage under targets and release using nuclease)(52,76) analyses of A_undiff_ and A_diff_ in adult testes. Using cell-number normalized spike-in controls, we quantitatively determined the genomic distributions of (i) PRC1-mediated H2AK119ub (by CUT&Tag), (ii) PRC2-mediated H3K27me3 (by CUT&Tag); and (iii) an active mark H3K4me3 (by CUT&RUN) in both isolated A_undiff_ and A_diff_ fractions (Figure 1D, Supplementary Figure S2). This spike-in approach enabled quantitative evaluation of changes in histone marks in spermatogonial differentiation and overcame the major limitation of previous epigenomic profiling studies that were not quantitative (37,38,45–47). We confirmed the high reproducibility between biological replicates (Supplementary Figure S2A), and these biological replicates were pooled for downstream analyses.

We found that H2AK119ub peaks are abundant throughout the genome in both A_undiff_ and A_diff_ (63,182 peaks in A_undiff_ and 47,409 peaks in A_diff_: Figure 1D, Supplementary Figure S2B), and are partially associated with H3K27me3, H3K4me3, or both (Supplementary Figure S2C, D, E, F). A notable change upon differentiation is a marked reduction of H3K27me3 enrichment and the number of genome-wide H3K27me3 peaks (Figure 1D). Among all H3K27me3 peaks detected across both A_undiff_ and A_diff_ (25,077 peaks), more than 50% (13,597 peaks) were identified as differential peaks (FDR<0.05), and most of these H3K27me3 peaks were lost upon differentiation (Figure 1E). Thus, our quantitative analysis uncovers a marked reduction of H3K27me3 as a characteristic of spermatogonial differentiation not noted in previous profiling studies (45,46).

Because PRCs bind gene promoters to regulate transcription(77–80), we next focused on the analysis of gene promoters. Among all Refseq gene promoters, 3,295 promoters showed differential enrichment of H3K27me3 from A_undiff_ to A_diff_, and 2,911 of them showed a significant loss of H3K27me3 (Figure 2A). To examine the relationship between chromatin state and gene regulation, we analyzed the enrichment of these marks at promoter regions of the differentially expressed genes (DEGs) during spermatogonial differentiation (identified in Figure 1C). Upregulated genes in A_diff_ were associated with a significant loss of H3K27me3 and a modest gain of H3K4me3 while maintaining stable H2AK119ub levels (Figure 2B, C). In contrast, downregulated genes in A_diff_ were associated with a modest gain of H2AK119ub on promoters without changes in H3K27me3 and H3K4me3 (Figure 2B, C). These features were evident at each representative locus: at the *Kit* gene locus, which is significantly induced in A_diff_, H3K27me3 was lost while H2AK119ub was maintained at its promoter during spermatogonial differentiation (Figure 2D). In contrast, at the *Lin28a* gene locus, a stem cell marker which is repressed in A_diff_, we saw increased H2AK119ub enrichment without apparent changes in H3K27me3 and H3K4me3 during spermatogonial differentiation (Figure 2E). The gain of H2AK119ub is in line with the recently reported function of a canonical PRC1(cPRC1) component CBX2 in spermatogonia (81).

**Figure 2.**
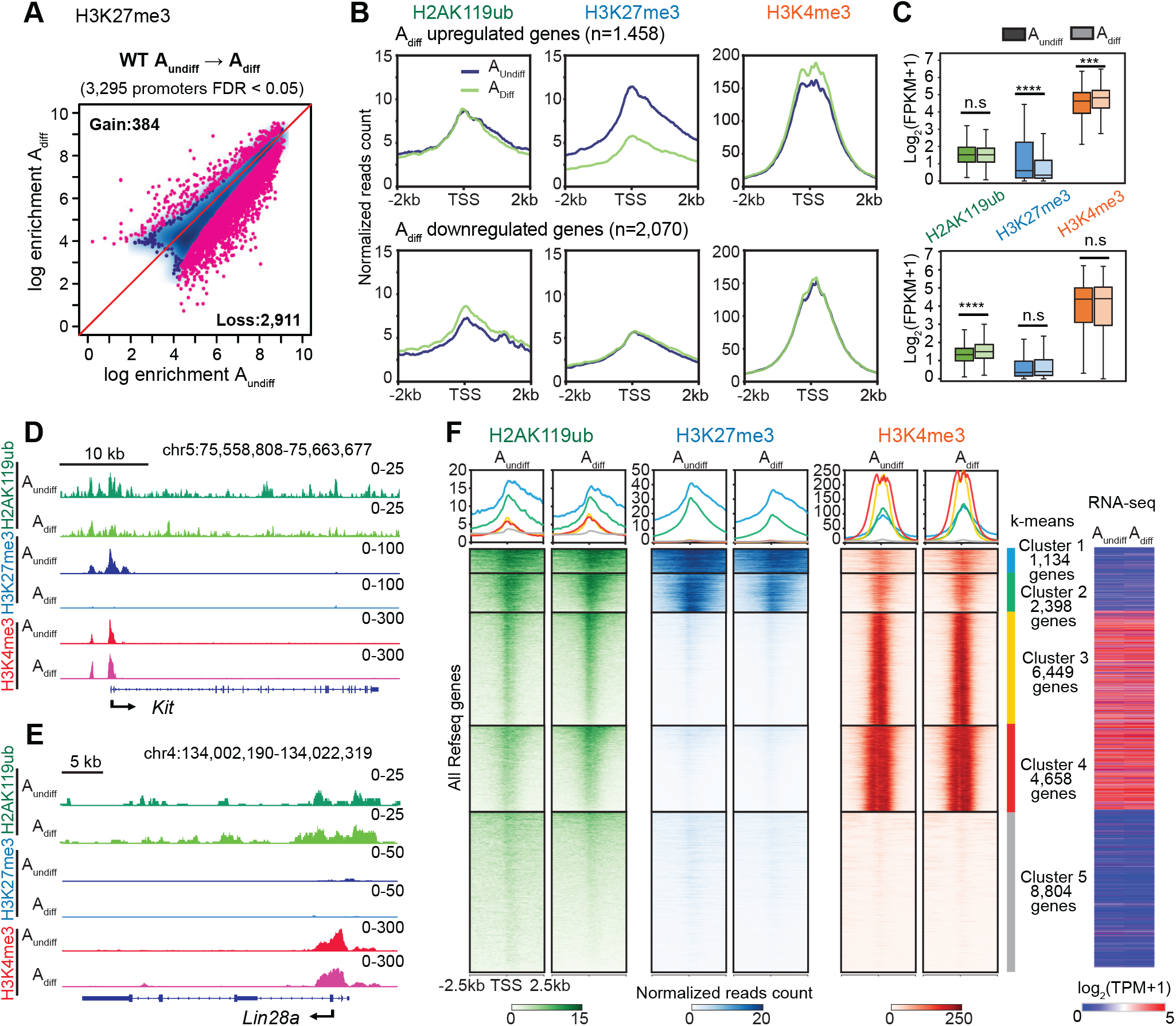
Spermatogonial differentiation is accompanied by a global loss of H3K27me3. (A) Scatter plot showing the comparison of H3K27me3 enrichment at promoters (TSS ± 2 kb) between A_undiff_ and A_diff_ in adult WT mice. Red dots represent genes with differential H3K27me3 enrichment at promoters (FDR < 0.05). (B) Average tag density plots showing H2AK119ub, H3K27me3, and H3K4me3 enrichment at promoter regions (TSS ± 2 kb) of DEGs during spermatogonial differentiation in adult WT mice (defined in Figure 1C). (C) Box-and-whisker plots showing H2AK119ub, H3K27me3, and H3K4me3 enrichment around TSSs in the corresponding gene groups shown in B. Central bars represent medians, the boxes encompass 50% of data points, and the whiskers include 90% of data points. n.s, not significant, *** P < 0.001, **** P < 5.1 × 10^-19^; two-tailed unpaired t-tests. (D and E) Track views of the *Kit* and *Lin28a* gene loci showing differential H2AK119ub, H3K27me3, and H3K4me3 occupancy in WT A_undiff_ and A_diff_. Data ranges in the upper right represent spike-in normalized read counts from combined replicates. (F) Heatmaps and average tag density plots of k-means clusters showing H2AK119ub, H3K27me3, and H3K4me3 enrichment at promoter regions (TSS ± 2.5 kb) in WT A_undiff_ and A_diff_. The color keys represent signal intensity, and the numbers represent spike-in normalized reads counts. Right: Heat map showing RNA-seq expression (log2-transformed TPM) of corresponding k-means clusters.

To capture genome-wide changes of these marks at promoters, we performed a k-means clustering analysis. In agreement with its genome-wide abundance, H2AK119ub was present on a broad range of promoters in clusters 1-4 (Figure 2F). H2AK119ub and H3K27me3 were similarly abundant in Cluster 1 and 2, which was also marked by weak H3K4me3 (Figure 2F). Cluster 1-2 genes are common targets of various Polycomb proteins (SUZ12, RNF2, MTF2, JARID2, and EZH2) in embryonic stem cells (ESCs; Supplementary Figure S3A) and are associated with transcription and developmental processes (Supplementary Figure S3B). Cluster 2 genes showed a marked reduction in H3K27me3 levels upon spermatogonial differentiation (Figure 2F). The presence of both H3K27me3 and H3K4me3 are characteristic of bivalent promoters, a chromatin context thought to maintain the potential for expression of developmental regulator genes in the germline (37,47,82,83). Thus, spermatogonial differentiation is accompanied by a reduction of H3K27me3 in a subset of bivalent promoters.

Notably, H2AK119ub was also enriched at promoters of active genes marked by high H3K4me3 level (Cluster 3 and 4) (Figure 2F), suggesting that H2AK119ub has a broader role than H3K27me3 in transcriptional regulation. These Cluster 3 and 4 genes are targets of various transcription factors, such as FOXO3, MYC, E2F1, and YY1 (Supplementary Figure S3A), and function in a broad range of biological processes, including RNA processing, DNA metabolic process, DNA repair, and cell cycle (Supplementary Figure S3B), which are constitutively active during spermatogonial differentiation (Figure 2F).

Together, these results demonstrate that H3K27me3 reduction is a prominent change that occurs during spermatogonial differentiation, raising the possibility that H3K27me3 shields the undifferentiated state of adult A_undiff_ to antagonize commitment to differentiation.

### PRC1 safeguards A_undiff_ from over-proliferation and differentiation

PRC1 and PRC2 are both required for the maintenance of undifferentiated spermatogonia (30,31). However, how PRCs function in steady-state spermatogenesis to sustain male fertility in the long term remains unknown. To elucidate this, we evaluated how the loss of PRC1 function affects steady-state spermatogenesis using a previously established conditional knockout mouse model (30,84). In brief, PRC1 function was eliminated after embryonic day 15 (E15) using *Ddx4*-Cre (50) to inactivate the E3 ubiquitin ligase RNF2 in a background lacking its partially redundant paralog, RING1 (RING1A) (30,48,84–86). These PRC1 conditional knockout (PRC1cKO) mice show nearly complete depletion of H2AK119ub in germ cells (30,84), representing a highly efficient loss-of-function PRC1 model.

Although PRC1 inactivation leads to gradual loss of the stem cell population and depletion of differentiated germ cells in postnatal spermatogenesis (30), A_undiff_ (detected based on the expression of PLZF, a marker for undifferentiated spermatogonia) were still present at 2 mo in PRC1cKO males (Figure 3A). By focusing on the analysis of PRC1cKO A_undiff_ at 2 mo, we sought to determine the role of PRC1 in steady-state spermatogenesis during which A_undiff_ slowly divide to maintain the stem cell pool. A previous study estimated that the doubling time of A_undiff_ is 3.9 days (93.6h), which becomes shorter once A_undiff_ commit to spermatogonial differentiation (28.5-34hr) (87). Therefore, we first evaluated the proportion of A_undiff_ with an active cell cycle at 2 mo by immunostaining with Ki67, a proliferation marker present in G1, S, G2, and M, but absent in G0 (88). We compared PRC1cKO males to littermate control mice (PRC1ctrl) which show normal spermatogenesis despite being *Ring1*^-/-^ (30). Consistent with the slow cycling of A_undiff_, only 17.4% of A_undiff_ in PRC1ctrl mice were Ki67^+^ (Figure 3A, B). However, 70.4% of A_undiff_ in the PRC1cKO testes were Ki67^+^, a ∼3-fold increase compared to PRC1ctrl males (Figure 3A, B), suggesting that PRC1cKO A_undiff_ are highly proliferative at 2 mo. However, the proportion of Ki67^+^ A_undiff_ was comparable between PRC1cKO and PRC1ctrl in juvenile testes at postnatal day 7 (P7) (30). Thus, the increased proliferation of A_undiff_ is a unique feature of steady-state spermatogenesis in PRC1cKO.

**Figure 3.**
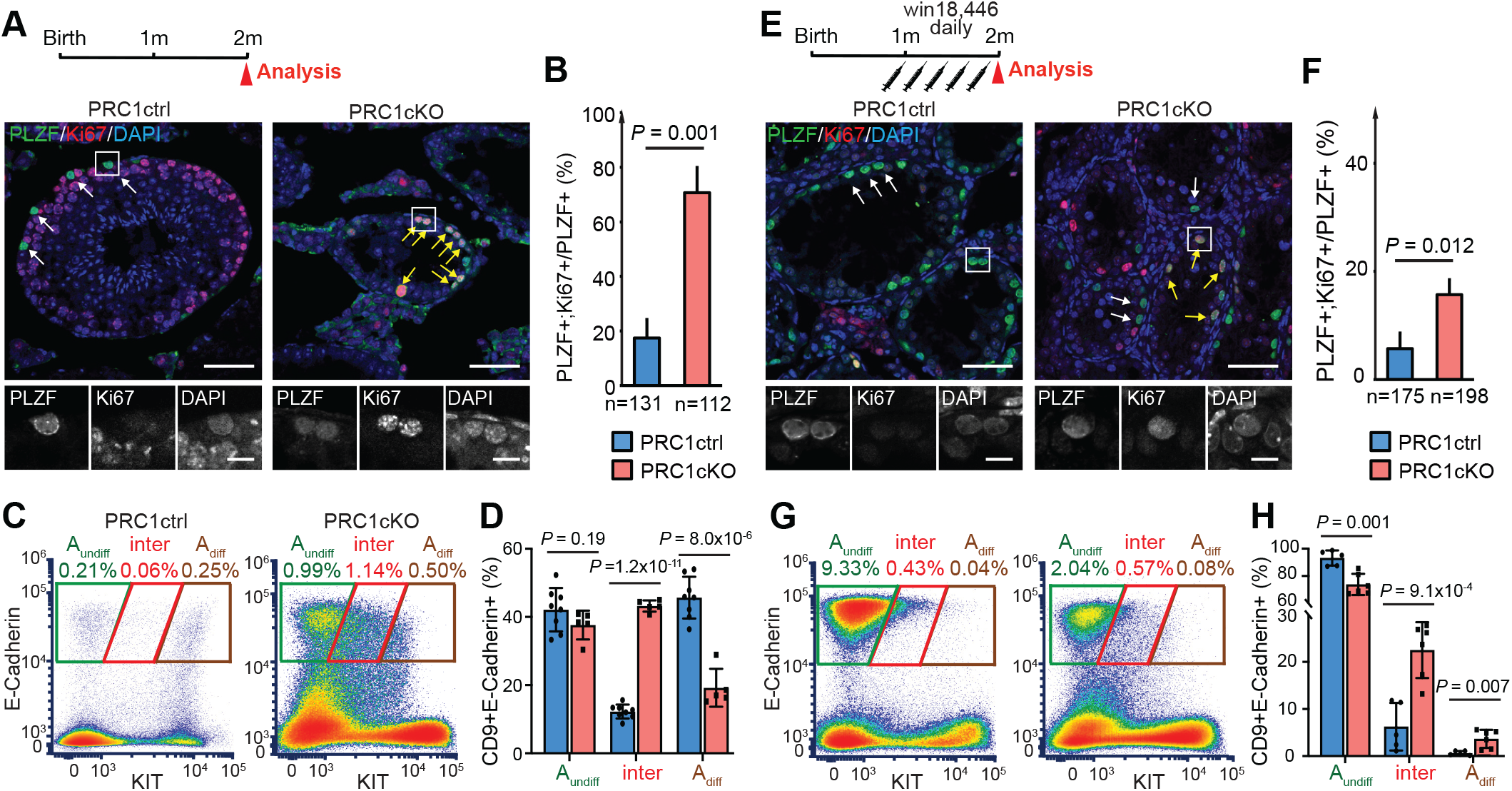
PRC1 deficiency causes over-proliferation of A_undiff_. (A and E) Schematics of the experimental workflow and immunostaining of testicular sections in PRC1cKO and a control littermate at 2 mo (A) or after win18,446 treatment (E). Ki67 (red) is a proliferation marker, and PLZF (green) is an undifferentiated spermatogonia marker. White arrows indicate the quiescent A_undiff_, and yellow arrows indicate A_undiff_ in the active cell cycle. Scale bars, 50 μm (10 μm in the boxed areas). (B and F) Quantitative analysis of immunostaining (A) and (E). Percentages representing numbers of PLZF^+^; Ki67^+^ cells per number of PLZF^+^ cells. n indicates the number of cells used for analysis. Data are represented as mean ± SD. Two-tailed unpaired Student’s t-tests. Three independent biological replicates were analyzed for each genotype. (C and G) Flow Cytometry profiles of all CD9^+^ cells from adult mouse testes in PRC1ctrl and PRC1cKO mice at 2 mo (C) or after win18,446 treatment (G). Percentages of each fraction relative to total cells are indicated. At least three independent biological replicates were analyzed for each genotype and representative profiles are shown. (D and H) Quantitative analysis of Flow Cytometry data (C) and (G). Percentages of each spermatogonial subpopulation per total CD9^+^E-Cadherin^+^ spermatogonia populations. Data are represented as mean ± SD. Two-tailed unpaired Student’s t-tests. Individual data points are also plotted.

To examine the cellular states of spermatogonia populations, we performed flow cytometry analyses of PRC1ctrl and PRC1cKO testicular cells at 2 mo. Like WT (Supplementary Figure S1), the CD9^+^E-Cadherin^+^ spermatogonia in PRC1ctrls consisted of two distinct populations (A_undiff_ and A_diff_; Figure 3C). By contrast, PRC1cKO spermatogonia did not show a clear separation between these two populations. Instead, intermediate state spermatogonia, that are transitioning to A_diff_, were abundant (CD9^+^E-Cadherin^+^ KIT^medium^ cells, abbreviated as “inter”, Figure 3C). 43.2% of total CD9^+^E-Cadherin^+^ spermatogonia in PRC1cKO males were transitioning, significantly higher than the 12.2% present in PRC1ctrls (Figure 3D). These results raise the possibility that PRC1 has cell-intrinsic roles in safeguarding A_undiff_ from over-proliferation and suppressing precocious commitment to differentiation.

Nevertheless, an alternative possibility is that the effects are cell-extrinsic: the above phenotypes could be indirectly caused by depletion of differentiated germ cells in PRC1cKO (Figure 3A) (30). As a result, A_undiff_ would become overcommitted to differentiation/proliferation in order to repopulate the depleted tubules. To distinguish between a cell-intrinsic and a cell-extrinsic effect resulting from PRC1 deletion, we blocked spermatogonial differentiation using Win18,446, an inhibitor of the enzyme aldehyde dehydrogenase 1a2 (ALDH1a2). This enzyme catalyzes endogenous RA biosynthesis from retinol (vitamin A) thus promoting RA-signaling which is essential for spermatogonial differentiation and meiosis initiation (89–92).

We reasoned that treatment with Win18,446 would block differentiation and enable evaluation of A_undiff_ without the influence of extrinsic differentiation signals. Pubertal PRC1ctrl and PRC1cKO males (1-month-old mice) were treated with Win18,446 injection daily until they fully matured at 2 months. After the treatment, both PRC1ctrl and PRC1cKO testes contained “empty” tubules devoid of differentiated germ cells, and only PLZF^+^ A_undiff_ were present, confirming the successful blockage of spermatogenesis after treatment with Win18,446 (Figure 3E). Due to RA-signaling inhibition, the proliferation rates of A_undiff_ were largely decreased in both PRC1ctrl and PRC1cKO, but PRC1cKO A_undiff_ were more susceptible to the RA-signaling inhibition (3.1-fold change from 17.4% to 5.7% in PRC1ctrl; versus 4.5-fold from 70.7% to 15.7% in PRC1cKO; Figure 3B, F). Overall, these results suggest that the RA-mediated differentiation cue extrinsically promotes the proliferation of A_undiff,_ and the effect of RA is more pronounced in the absence of differentiated germ cells in PRC1cKO mice. However, the proliferative activity of A_undiff_ in the PRC1cKO was still significantly higher than in PRC1ctrl testes (15.7% versus 5.7% of all A_undiff_; Figure 3F), suggesting that PRC1 cell-intrinsically suppresses the proliferation of A_undiff_. In addition, enrichment of intermediate state spermatogonia was still observed after treatment with Win18,446 (Figure 3G, H). Altogether, we conclude that PRC1 cell-intrinsically safeguards A_undiff_ from over-proliferation/differentiation.

### PRC1 suppresses the spermatogonial differentiation program in A_undiff_

To determine the function of PRC1 in A_undiff_, using the FACS method described above, we isolated A_undiff_ from 2-month-old PRC1ctrl and PRC1cKO testes (identical E-Cadherin^+^CD9^high^KIT^−^ population) and performed bulk RNA-seq (Supplementary Figure S4A). We found that, in PRC1cKO, 230 genes were significantly upregulated, and 303 genes were significantly downregulated compared to PRC1ctrls (Figure 4A and Supplementary Data 2). Notably, upregulated genes included those known to be related to spermatogonial differentiation (*Kit, Esx1,* and *Crabp1*) as well as meiotic processes (*Prdm9, M1ap,* and *Meioc)*, whereas genes required for A_undiff_ maintenance (*Sall4, Lin28a, Nanos3,* and *Sall1*) were downregulated (Figure 4A). We further performed an RNA-seq analysis of A_undiff_ after treatment with Win18,446, to minimize cell-extrinsic effects (Figure 4B; Supplementary Figure S4B). Under this condition, more genes were misregulated in PRC1cKO compared to PRC1ctrl (570 genes upregulated and 836 genes downregulated), and, of note, spermatogonial differentiation genes (e.g., *Kit, Dmrt1, Dmrtb1, Esx1,* and *Tcea3*) were still significantly upregulated after the treatment (Figure 4B, Supplementary Data 2).

**Figure 4.**
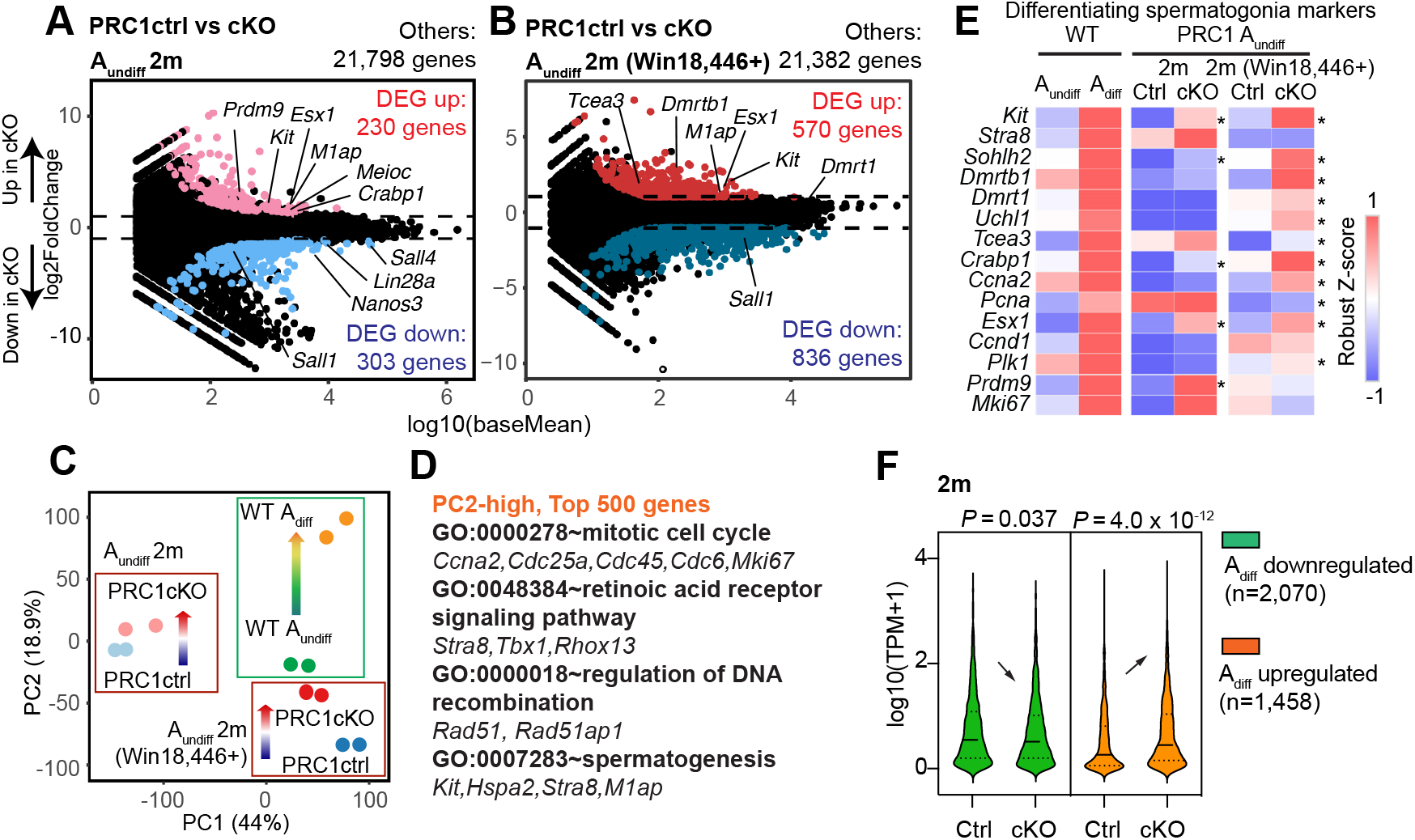
PRC1 deficiency in A_undiff_ leads to precocious activation of the spermatogonial differentiation program. (A and B) Comparison of PRC1ctrl and PRC1cKO A_undiff_ transcriptomes at 2 mo (A) or after win18,446 treatment (B). Two independent biological replicates were examined for RNA-seq. All genes with False Discovery Rate (FDR) values are plotted. Differentially expressed genes (DEGs: Log2FoldChange >1, FDR< 0.05, TPM >1) are colored (red: upregulated in PRC1cKO A_undiff_; blue: downregulated in PRC1cKO A_undiff_), and numbers are shown. (C) PCA plot showing transcriptome relationships between A_undiff_ and A_diff_ in WT, and A_undiff_ between PRC1ctrl and PRC1cKO at 2 mo with or without Win18,446. Arrows indicate increased PC2 loading. (D) GO enrichments and representative genes contributing to high PC2 loading (top 500 genes) in PCA (C) are listed. (E) Heatmap showing gene expression of representative differentiating spermatogonia markers in PRC1ctrl and PRC1cKO A_undiff_ at 2 mo with or without Win18,446 treatment, as well as in A_undiff_ and A_diff_ of WT. *, FDR< 0.05. (F) Violin plots showing the expression levels of WT spermatogonia DEG groups detected in Figure 1C in PRC1ctrl and PRC1cKO A_undiff_ at 2 mo. Two-tailed unpaired t-tests.

It is worth noting that the *Cdkn2a* gene, which encodes two major cell cycle inhibitors (p16 (INK4A) and p14 (ARF)) and is a major Polycomb target for suppression through which Polycomb promotes proliferation in many other tissue stem cells and somatic cells, was not upregulated in the PRC1cKO A_undiff_ (Supplementary Figure S4C), consistent with previous studies using Polycomb mutants in spermatogenesis (30,31). Therefore, *Cdkn2a* is not involved in the PRC1-mediated regulation of cell cycle in spermatogonia. GO analysis revealed that upregulated genes in both PRC1cKO A_undiff_ samples were enriched for cell-cell adhesion and cell fate commitment, processes associated with development/morphogenesis (Supplementary Figure S4D, E). In turn, downregulated genes were over-represented with diverse molecular functions, including stem cell population maintenance, spermatogenesis, and chromatin organization implying compromised spermatogenic programs in PRC1cKO A_undiff_ (Supplementary Figure S4D, E).

To clarify the role of PRC1 in spermatogonial differentiation, we compared all RNA-seq data in this study by Principal-component analysis (PCA) to capture overall transcriptomic changes among all pairs of samples (Figure 4C). A transcriptome change during spermatogonial differentiation in WT (from WT A_undiff_ to WT A_diff_) was accompanied by increased PC2 loading. Similar directional changes (PC2 increase) were observed in PRC1ctrl A_undiff_ and PRC1cKO A_undiff_ both in non-treated and Win18,446-treated conditions. Notably, the genes contributing to the positive PC2 values were enriched with those upregulated during spermatogonial differentiation, involved in mitotic cell cycle, retinoic acid receptor signaling pathway, and spermatogenesis. Examples include *Ccna2, Mki67, Stra8,* and *Kit* (Figure 4D).

To validate this observation, we next examined the expression level of previously identified genes that are upregulated during spermatogonial differentiation (75). These differentiating spermatogonia marker genes tended to show increased expression levels in the A_undiff_ of PRC1cKO males both in non-treated and Win18,446-treated conditions compared to PRC1ctrls (Figure 4E). We also tested how the DEGs detected in WT spermatogonial differentiation (shown in Figure 1C) are regulated by PRC1. Genes upregulated in WT A_diff_ showed significantly higher expression in the A_undiff_ of PRC1cKO than PRC1ctrl (Figure 4F), suggesting they were precociously activated in mutant A_undiff_. By contrast, genes active in WT A_undiff_ showed decreased expression in PRC1cKO A_undiff_ compared to PRC1ctrls.

Collectively, genes linked to spermatogonial differentiation were precociously activated in PRC1cKO A_undiff_ whereas those linked to A_undiff_ identity were present at decreased levels. Therefore, we conclude that PRC1 shields the undifferentiated state of adult A_undiff_ by suppressing the spermatogonial differentiation program.

### Removal of PRC1 impairs the global deposition of H3K27me3 in A_undiff_

To determine the mechanism by which PRC1 regulates spermatogonial differentiation at the chromatin level, we applied quantitative CUT&RUN to profile the genomic distributions of H3K27me3 and H3K4me3 in FACS-purified A_undiff_ from PRC1ctrl and PRC1cKO at 2 months of age (Figure 5A and Supplementary Figure S5A). Since our data show that H3K27me3 levels are a major difference between A_undiff_ and A_diff_ in WT (Figure 2), we sought to determine how PRC1 regulates PRC2-mediated H3K27me3. Remarkably, the H3K27me3 landscape was greatly impaired in PRC1cKO A_undiff_ with a dramatic reduction of both peak numbers and intensity (Supplementary Figure S5B, C, D). H3K27me3 peak distribution changed in various genomic regions, including gene bodies and intergenic regions (Supplementary Figure S5C), and a global loss of H3K27me3 at gene promoters was observed in PRC1cKO A_undiff_ (Figure 5A). The active mark, H3K4me3, was instead slightly increased at gene promoters (Figure 5A). These features are exemplified by track views of the *Kit, Stra8,* and *Tcea3* loci (Figure 5B), where H3K27me3 nearly completely disappeared at promoters, consistent with transcriptional de-repression. These results suggest that PRC1 regulates the distribution of H3K27me3 and counteracts H3K4me3 in A_undiff_. After treatment with Win18,446, genome-wide H3K27me3 deposition was still largely impaired in PRC1cKO A_undiff_ (Supplementary Figure S5B, C, D), and there was a prominent loss of H3K27me3 at gene promoters (Figure 5A), indicating that the PRC1 regulation of H3K27me3 is a cell-intrinsic feature of A_undiff_. By contrast, after treatment with Win18,446, H3K4me3 slightly decreased at promoters of PRC1cKO A_undiff_, as opposed to the slight increase in non-treated PRC1cKO A_undiff_, suggesting that H3K4me3 changes were secondary consequences of PRC1 deletion.

**Figure 5.**
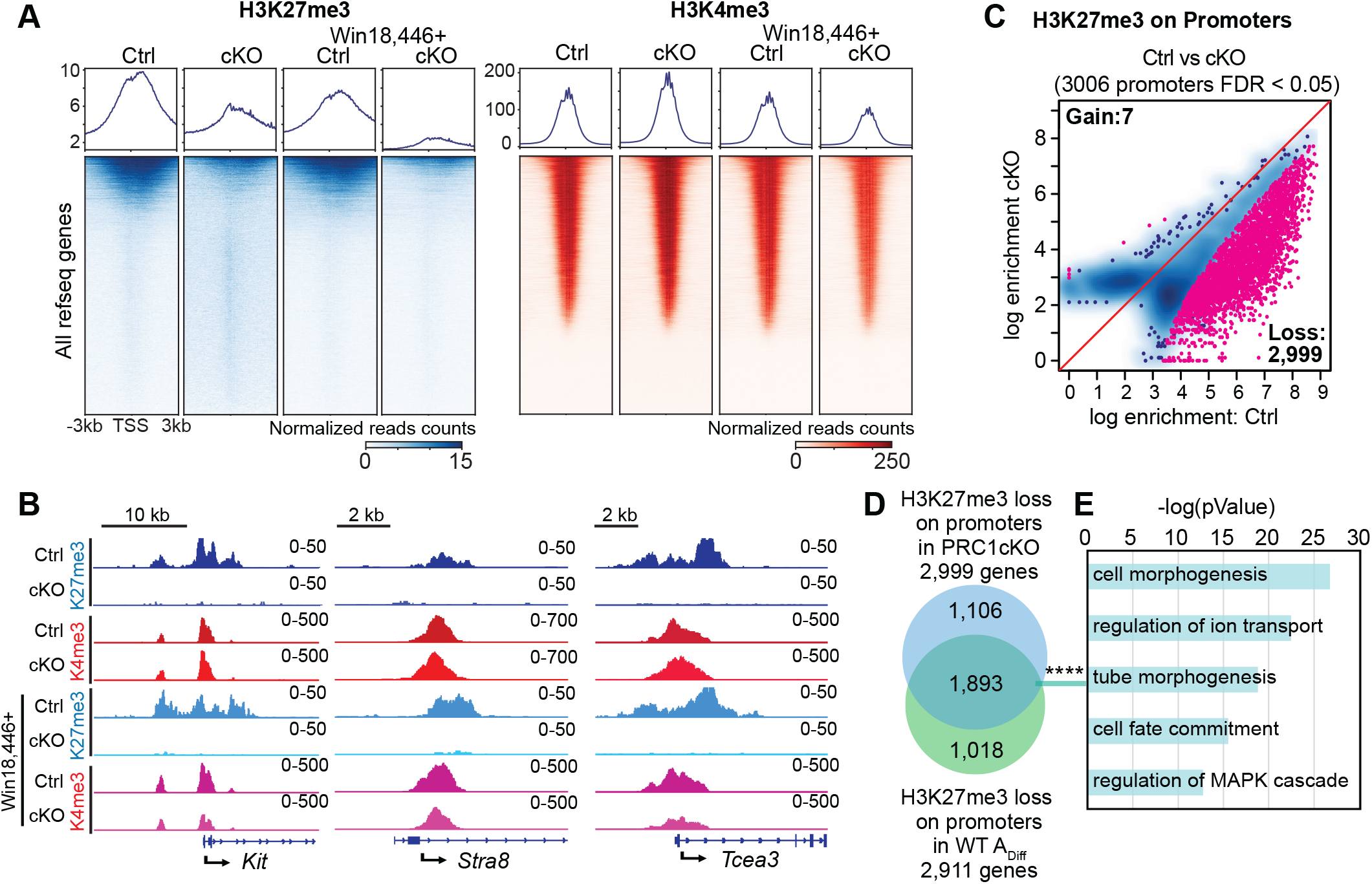
Removal of PRC1 impairs the global deposition of H3K27me3 in Aundiff genomes. (A) Heatmaps and average tag density plots showing H3K27me3 and H3K4me3 enrichment at promoter regions (TSS ± 3 kb) in PRC1ctrl and PRC1cKO A_undiff_ at 2 mo with or without Win18,446 treatment. The bars below the heatmaps represent signal intensity, and the numbers represent spike-in normalized reads counts. (B) Track views of the *Kit*, *Stra8* and *Tcea3* gene loci showing H3K27me3 and H3K4me3 CUT&RUN peaks in A_undiff_ of indicated genotypes. Data ranges are shown in the upper right. (C) Scatter plot showing comparison of H3K27me3 enrichment at promoters (TSS ± 2 kb) between PRC1ctrl and PRC1cKO A_undiff_ at 2 mo. Red dots represent genes with differential H3K27me3 enrichment (FDR < 0.05). (D) Overlap between H3K27me3-lost promoters in PRC1cKO A_undiff_ and H3K27me3-lost promoters in WT A_diff_. ****P = 2.8 x 10^-1207^, two-sided hypergeometric test. (E) Key GO enrichments in the overlap genes detected in D.

We further identified 3,006 promoters with significant changes in H3K27me3 occupancy in PRC1cKO A_undiff_, and 2,999 of them lost H3K27me3 (Figure 5C), resembling changes that occur during WT spermatogonial differentiation. Remarkably, genes showing H3K27me3 loss at promoters in PRC1cKO A_undiff_ significantly overlapped with genes that lost H3K27me3 in WT A_diff_ (Figure 5D); 1,893 overlapping genes included those involved in cell/tube morphogenesis, cell fate commitment and MAPK signaling (Figure 5E, Supplementary Data 3). A similar overlap was observed in PRC1cKO after treatment with Win18,446 (Supplementary Figure S6A, B, C, Supplementary Data 3). These results suggest that the overall chromatin state of PRC1cKO A_undiff_ resembles that of WT A_diff_, raising the possibility that PRC1 regulates PRC2-mediated H3K27me3 to shield the chromatin state in A_undiff_ and prevent precocious differentiation.

### PRC1 suppresses precocious spermatogonial differentiation and ectopic somatic program in A_undiff_

Because PcG proteins regulate a broad range of developmental genes that are critical for pattern formation and cell fate specification in development, we next sought to determine how PRC1 regulates developmental genes in A_undiff_. To this end, we used a list of canonical PcG target genes that are known to be suppressed to maintain developmental potential in ESCs (93). We reasoned that PRC1 shields the undifferentiated state by defining an A_undiff_-specific gene expression program. Indeed, canonical PcG target genes largely overlapped with genes that lose H3K27me3 at promoters in WT spermatogonial differentiation and genes that carry PRC1-dependent H3K27me3 at promoters (Supplementary Figure S6D, E).

We first categorized the canonical PcG target genes into three major groups based on their expression dynamics during spermatogonial differentiation from A_undiff_ to A_diff_ in WT: (1) 608 upregulated PcG target genes (PcG up); (2) 618 downregulated PcG target genes (PcG down); (3) 2,373 PcG targets that did not show significant changes (PcG stable: Figure 6A). A notable feature was decreased H3K27me3 at entire regions of “PcG up” genes (TSSs, gene bodies, transcription termination sites, TESs) during WT spermatogonial differentiation (Figure 6B). On these “PcG up” genes, H3K27me3 was largely PRC1-dependent as shown in PRC1cKO A_undiff_ with or without Win18,446 (Figure 6C), indicating that the loss of PRC1-dependent H3K27me3 occurred upon spermatogonial differentiation. Quantification of H3K27me3 at promoters further confirmed this conclusion (Figure 6D). These “PcG up” genes were upregulated in PRC1cKO A_undiff_ (Figure 6E). Therefore, in A_undiff_, PRC1 prevents precocious activation of developmental genes linked to spermatogonial differentiation.

**Figure 6.**
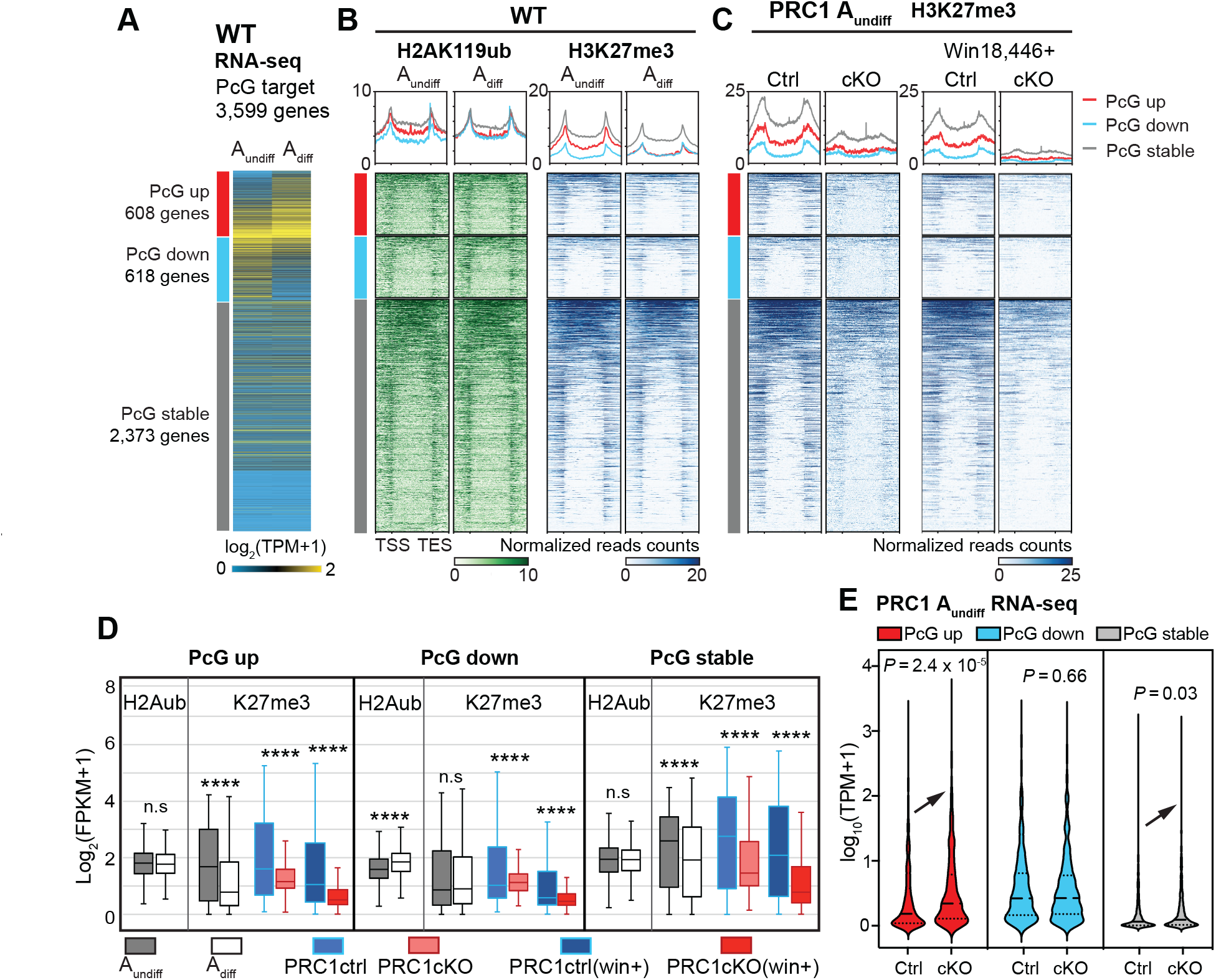
Polycomb shields the A^undiff^-specific developmental program in adult spermatogonia. (A) Heatmap showing RNA-seq expression (log2-transformed TPM) of corresponding Polycomb-targets groups. (B and C) Heatmaps and average tag density plots of H2AK119ub, H3K27me3, and H3K4me3 enrichment at genic regions (TSS-TES ± 2 kb) on different Polycomb-target groups in WT A^undiff^ and A^diff^ (B) or PRC1 A^undiff^ of indicated genotypes (C). The bars below the heatmaps represent signal intensity, the numbers represent spike-in normalized reads counts. (D) Box-and-whisker plots showing H2AK119ub and H3K27me3 enrichment at promoters (TSS ± 2 kb, log2-transformed FPKM) in the corresponding gene groups shown in B and C. Central bars represent medians, the boxes encompass 50% of the data points, and the whiskers indicate 90% of the data points. n.s, not significant, **** P < 4.1 x 10^-6^; two-tailed unpaired t-tests. (E) Violin plots showing the expression level of different Polycomb-targets groups in 2 mo A^undiff^ of PRC1ctrl and PRC1cKO.

These results also suggest that PRC1 plays a major role in restricting transcription of a large number of PcG targets through the regulation of PRC2 during spermatogonial differentiation. In addition, decreased H3K27me3 was also observed on “PcG stable” genes during spermatogonial differentiation (Figure 6B), and, in A_undiff_, this is largely PRC1-dependent (Figure 6C, D). The “PcG stable” genes were ectopically upregulated in PRC1cKO A_undiff_ (Figure 6E). Therefore, in A_undiff_, PRC1 suppresses ectopic activation of developmental genes that are required for somatic development. Furthermore, increased H2AK119ub was observed on PcG down genes (Figure 6D), suggesting that PRC1 directly represses developmental genes during spermatogonial differentiation. Although this PRC1-dependent regulation of H3K27me3 was consistent regardless of Win18,446 treatment, H3K4me3 showed distinct changes in the presence or absence of Win18,446 (Supplementary Figure S6F, G), further demonstrating an indirect role for PRC1 in H3K4me3 regulation.

Misregulated gene expression at the *Hoxa* cluster, which includes classic Polycomb-targeted developmental genes, exemplifies the changes observed in PRC1cKO A_undiff_ (Supplementary Figure S7). These *Hoxa* genes were regulated differently during WT spermatogonial differentiation, e.g., the *Hoxa1, Hoxa2,* and *Hoxa3* gene loci substantially lost H3K27me3, resulting in transcriptional upregulation; however, the *Hoxa9, Hoxa10,* and *Hoxa11* genes gained H2AK119ub and were downregulated. In PRC1cKO A_undiff_, H3K27me3 was depleted across the entire *Hoxa* cluster, leading to ectopic de-repression of *Hoxa* genes. These results demonstrate that PRC1 has two major functions in A_undiff_: preventing precocious activation of developmental genes that are activated later during spermatogonial differentiation and suppressing ectopic expression of the somatic program. Together these functions shield the undifferentiated state by defining an A_undiff_-specific gene expression program.

### PRC1 is a negative regulator of the cell cycle in A_undiff_

Because we found increased proliferation of the PRC1cKO A_undiff_ that leads to A_undiff_ exhaustion, we next examined the PRC1-dependent regulation of cell cycle-related genes. We found that H2AK119ub accumulated on genes that positively regulate the cell cycle, e.g., cyclins, including *Ccna1, Ccnd1, Ccne1,* and *Ccne2*; the cyclin-dependent kinases, *Cdk2* and *Cdk5*; cell division regulators, *Cdc6* and *Cdc25a*; as well as the cell cycle inhibitor, *Cdkn1b*; suggesting direct regulation of these genes by PRC1 (Figure 7A, Supplementary Figure S8A, B). Consistent with the over-proliferation phenotype of PRC1cKO A_undiff_, increased expression of these pro-cell cycle regulators and decreased *Cdkn1b* expression in PRC1cKO A_undiff_ were observed (Supplementary Figure S8A, B). Among these genes, H3K27me3 only accumulated on the *Ccna1* gene, indicating that PRC1 regulates the other genes independent of PRC2 (Figure 7A). Together with the over-proliferation phenotype of PRC1cKO cells, these results demonstrate that PRC1 is a negative regulator of the cell cycle in A_undiff_ that maintains the characteristic slow cycling of A_undiff_. Therefore, the slow cycling of A_undiff_ is part of a PRC1-directed germline program that protects the undifferentiated state of adult A_undiff_ and regulates spermatogonial differentiation.

**Figure 7.**
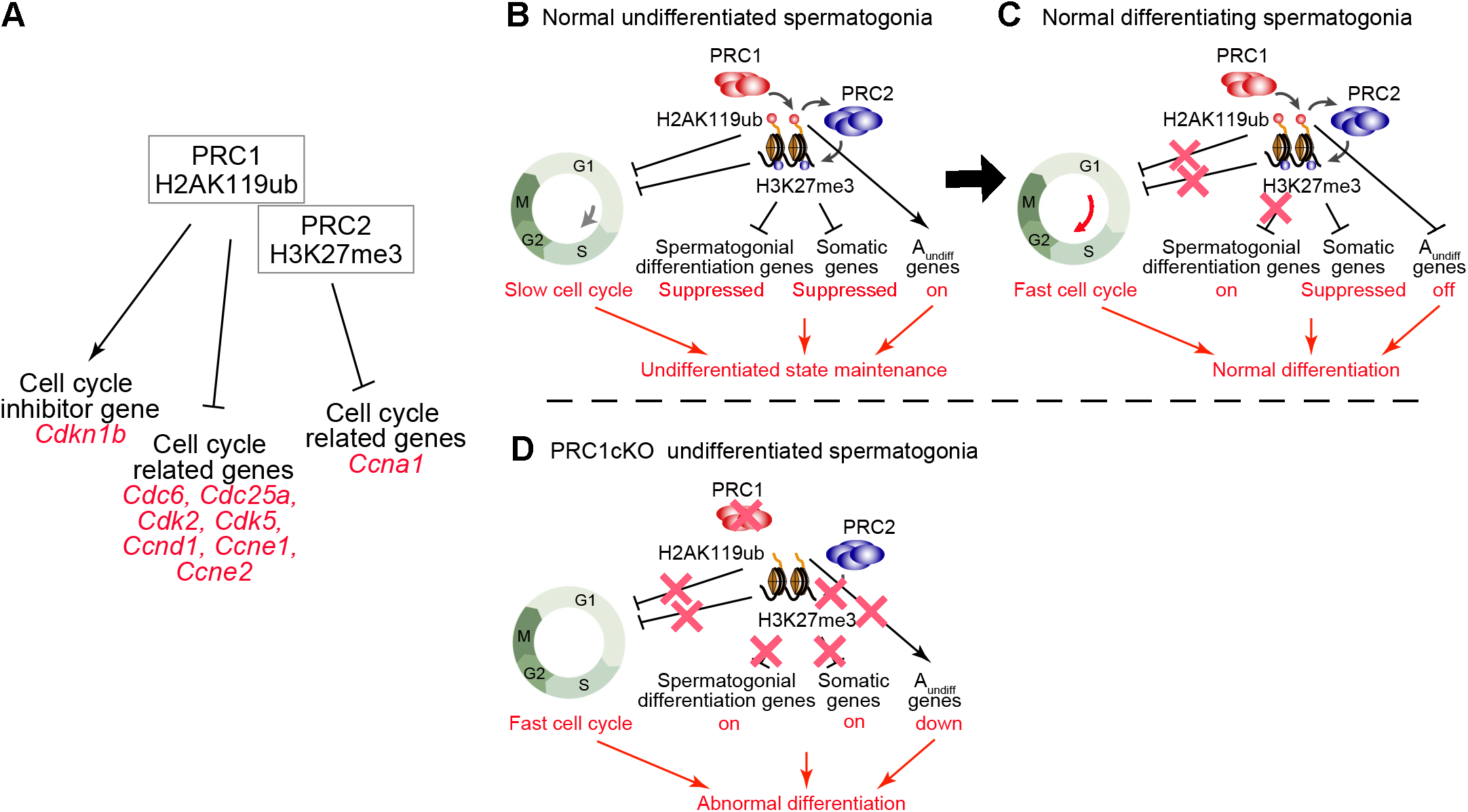
Models of Polycomb functions in adult A_undiff_ and spermatogonial differentiation. (A) Summary of cell cycle regulation by PRC1 and PRC2. (B, C and D) Models of Polycomb functions that shield the undifferentiated state of A_undiff_ (B), regulate spermatogonial differentiation (C), and defective spermatogonial differentiation in the absence of PRC1 (D).

## Discussion

In this study, we find that PRC1 plays a central role in defining the A_undiff_-specific developmental program and maintaining a long-lasting chromatin state in A_undiff_ during adult steady-state spermatogenesis. PRC1 shields the undifferentiated state of A_undiff_ from spermatogonial differentiation. Furthermore, PRC1 is a negative cell cycle regulator that maintains slow cycling of A_undiff_. In the absence of PRC1, cell proliferation increased in A_undiff_ both cell-intrinsically and cell-extrinsically, and A_undiff_ committed to abnormal differentiation. Our quantitative epigenomic analysis clarified the dynamics of chromatin changes during spermatogonial differentiation, an aspect not quantitatively evaluated in previous studies (37,38,45–47). Importantly, we show that H3K27me3 is a major epigenetic hallmark of A_undiff_ maintenance. Indeed, spermatogonial differentiation is accompanied by a global resolution of bivalent domains through the loss of H3K27me3 on genes required for spermatogonial differentiation. Our data argue that PRC1 directs PRC2-H3K27me3 deposition, thus supporting A_undiff_ maintenance (Figure 7B, C). Overall, we have determined a molecular logic underlying spermatogonial differentiation: the loss of H3K27me3 at promoters activates the differentiation program (Figure 7B, C).

In WT spermatogonial differentiation, an extensive loss of H3K27me3 at target promoters does not necessarily lead to massive gene de-repression in normal A_diff_, presumably due to constant genome-wide occupancy of H2AK119ub. Hence, H3K27me3 does not maintain constitutive repression at bivalent promoters alone; instead, H2AK119ub acts as an additional safety lock to ensure developmental gene regulation. Further, spermatogonial differentiation occurs with the suppression of A_undiff_ genes by PRC1 independent of PRC2. Without PRC1, both spermatogonial differentiation genes and somatic genes were ectopically activated, leading to abnormal differentiation (Figure 7D).

Our study demonstrates extensive direct regulation of PRC2-H3K27me3 by PRC1-H2AK119ub *in vivo*. Although early studies showed that cPRC1 acts downstream of PRC2-deposited H3K27me3 (94,95), recent studies have instead demonstrated that variant PRC1 (vPRC1) mediates H2AK119ub-dependent PRC2 recruitment (96–98). We reanalyzed recent cPRC1 CUT&RUN data in adult spermatogonia (81) and found that key cPRC1 components, including CBX2, PHC2, and BMI1, bind a small portion of PRC2-H3K27me3 target genes specifically in KIT^+^ differentiating spermatogonia (Supplementary Figure S9A). This analysis supports the hypothesis that vPRC1 plays a major role in regulating PRC2-H3K27me3 in A_undiff_. We also note that cPRC1 and vPRC1 are regulated differently during spermatogonial differentiation. Several genes encoding cPRC1 components, such as *Pcgf2*, *Cbx2/4*, and *Phc2* genes, show increased expression as spermatogonia move from the A_undiff_ to A_diff_ state (Supplementary Figure S9B). However, the regulation of vPRC1 seems complicated; while the expression of the *Pcgf3/5* genes (encoding the components of vPRC1.3 and 1.5) is increased, the expression of the *Pcgf6* gene (encoding the components of vPRC1.6) is decreased, raising the possibility that vPRC1 subcomplexes are reorganized upon spermatogonial differentiation. Further studies are needed to clarify the regulation of PRC1 subcomplexes in spermatogonial differentiation.

Major questions raised in this study are how H3K27me3 loss occurs at the onset of spermatogonial differentiation and what triggers this process. There are three possible mechanisms that could account for H3K27me3 loss. One is decreased PRC2 activity, the second is H3K27 demethylation, and the third is histone H3 replacement with a testis-specific histone variant, H3T (99). We found that the expression levels of PRC2 components are largely consistent; however, the H3K27 demethylase KDM6B/JMJD3 is significantly upregulated from A_undiff_ to A_diff_ (Supplementary Figure S9B). In line with this observation, a germline-specific deletion of KDM6B leads to increased and prolonged maintenance of undifferentiated spermatogonia (100). Therefore, we speculate the KDM6B actively demethylates H3K27me3 at specific target loci at the onset of spermatogonial differentiation. As for the trigger of H3K27me3 loss, we suspect that RA signaling plays a role. This is because our reanalysis of a previous study (101) suggests that RA receptor (RAR) and STRA8 bind to the promoters of the *Rnf2*, *Eed* (PRC2 component), and *Kdm6b* genes; this RAR binding increases in response to RA (Supplementary Figure S9C). These data suggest that the RA signaling may trigger the expression of the *Kdm6b* gene, leading to H3K27me3 loss as A_undiff_ differentiate. Intriguingly, the RA receptor gene loci (*Rara*, *Rarb*, *and Rarg)* and the *Stra8* gene locus are also targeted by PRC1/PRC2 (Supplementary Figure S9D), suggesting that the Polycomb system and RA-STRA8 signaling may form a regulatory loop to coordinate each other’s expression.

We note that H2AK119ub is pervasively enriched at highly active genes (Figure 2F; Cluster3-4) during spermatogonial differentiation, a phenomenon that has been reported in many other contexts, including growing oocytes (102), skin stem cells (103), and leukemic cells (104). Dual roles for PRC1 in gene activation and repression were also reported in juvenile spermatogonia (30). In adult spermatogonia, in a small portion of H2AK119ub/H3K4me3 co-enriched active genes, we found both upregulated and downregulated DEGs in WT A_diff_ or PRC1cKO A_undiff_ (Supplementary Figure S9E), supporting a mechanism where PRC1 fine-tunes gene repression and activation. However, it is unlikely that local H2AK119ub at active gene promoters plays a critical role in gene regulation because these genes are not greatly dysregulated in PRC1cKO A_undiff_.

Notably, these gene regulatory mechanisms are tightly coupled with cell cycle regulation by PRC1. This feature is unique to A_undiff_ because PRC1 positively regulates the cell cycle in somatic cells by suppressing the major cell cycle inhibitor *Cdkn2a* gene, and the loss of PcG-related functions leads to cell cycle inhibition (105,106). In contrast, a *Cdkn2a-*independent function of PRC1 regulates the slow cycling property of A_undiff_ prior to the increased proliferation observed upon spermatogonial differentiation. We found that PRC1 directly suppresses genes that activate the cell cycle and promotes expression of the cell cycle inhibitor *Cdkn1b*, thereby functioning as a negative regulator of the cell cycle in A_undiff_. In PRC1cKO A_undiff,_ increased expression of these pro-cell cycle regulators and decreased *Cdkn1b* expression likely account for their over-proliferation phenotype. Alternatively, increased expression of pro-apoptotic genes, e.g., *Bok, Bax,* and *Casp8,* resulting from loss of PRC1 activity at these loci could also contribute to the gradual loss of A_undiff_ in PRC1cKO testes (Supplementary Figure S8C).

Our study uncovers PRC1-directed mechanisms for the maintenance and differentiation of undifferentiated spermatogonia in adults. These newly defined Polycomb-regulated mechanisms are likely to be related to the unidirectional differentiation program of spermatogenesis, in which the chromatin states of progenitor cells are predetermined to direct spermatogenic differentiation (47,107). Similarly, PRC2-H3K27me3 functions in human pluripotent cells to shield the potency of stem cells (108,109); thus, a parallel mechanism may underlie stem cell maintenance in general. Given the persistent nature of H3K27me3 in the germline (37,45,47,110), PRC1-directed regulation of PRC2-H3K27me3 could be a pivotal mechanism to sustain germline potential and to recover totipotency and somatic gene regulation after fertilization.

## Data availability

RNA-seq data and CUT&RUN/Tag datasets were deposited in the Gene Expression Omnibus under accession no. GSE221944. Source data are provided in this paper.

## Code availability

Source code for all software and tools used in this study, with documentation, examples, and additional information, is available at the URLs listed above.

## Author contributions

M.H. and S.H.N. designed the study in discussion with S.Y. M.H., Y.H.Y., and S.M. performed experiments. M.H. and S.H.N. designed and interpreted the computational analyses. M.H. and T.N. developed the protocol to isolate spermatogonia. M.H., S.Y., and S.H.N. interpreted the results. M.H. and S.H.N. wrote the manuscript with critical feedback from all other authors. S.H.N. supervised the project.

## Acknowledgments

We thank members of the Namekawa lab, Richard Schultz and Neil Hunter, for discussion, Miguel Vidal and Haruhiko Koseki for sharing *Ring1^-/-^ and Rnf2^F/F^* mouse lines, Artem Barski for sharing the reagents.

