## Supplemental figures for "PRC1 directs PRC2-H3K27me3 deposition to shield adult spermatogonial stem cells from differentiation"

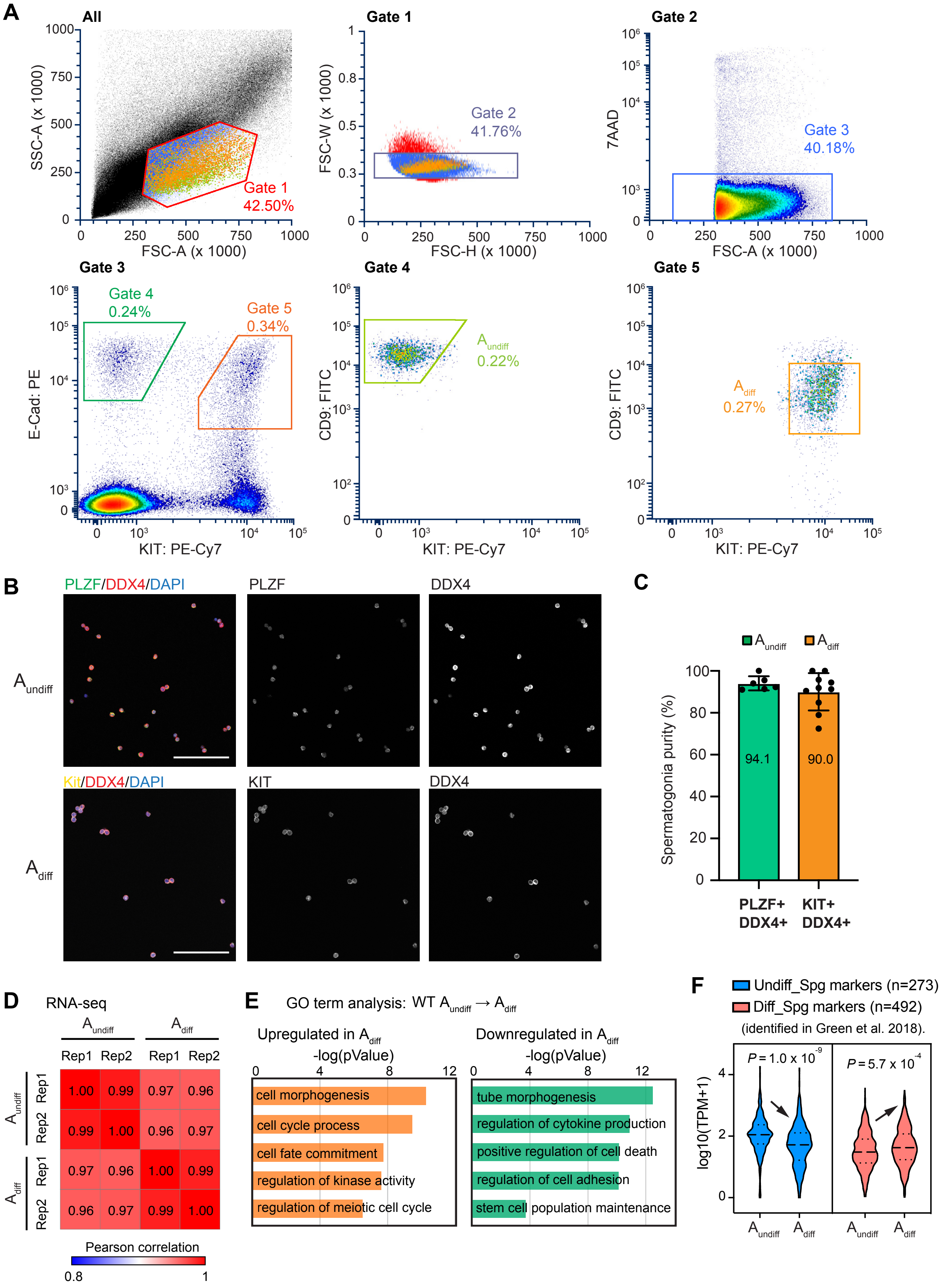

A
All
SSC-A (x 1000)
FSC-A (x 1000)
Gate 1
42.50%
Gate 1
FSC-W (x 1000)
FSC-H (x 1000)
Gate 2
41.76%
Gate 2
7AAD
FSC-A (x 1000)
Gate 3
40.18%
Gate 3
E-Cad: PE
KIT: PE-Cy7
Gate 4
0.24%
Gate 5
0.34%
Gate 4
CD9: FITC
KIT: PE-Cy7
A undiff
0.22%
Gate 5
CD9: FITC
KIT: PE-Cy7
A diff
0.27%
B
PLZF/DDX4/DAPI
PLZF
DDX4
A undiff
Kit/DDX4/DAPI
KIT
DDX4
A diff
C
A undiff
A diff
Spermatogonia purity (%)
94.1
90.0
PLZF+ DDX4+
KIT+ DDX4+
D
RNA-seq
A undiff
A diff
Rep1
Rep2
Rep1
Rep2
1.00 0.99 0.97 0.96
0.99 1.00 0.96 0.97
0.97 0.96 1.00 0.99
0.96 0.97 0.99 1.00
Pearson correlation
0.8
1
E
GO term analysis: WT A undiff → A diff
Upregulated in A diff
-log(pValue)
cell morphogenesis
cell cycle process
cell fate commitment
regulation of kinase activity
regulation of meiotic cell cycle
Downregulated in A diff
-log(pValue)
tube morphogenesis
regulation of cytokine production
positive regulation of cell death
regulation of cell adhesion
stem cell population maintenance
F
Undiff_Spg markers (n=273)
Diff_Spg markers (n=492)
(identified in Green et al. 2018).
P = 1.0 x 10-9
P = 5.7 x 10-4
log10(TPM+1)
A undiff
A diff
A undiff
A diff

**Figure S1 Gene expression profiles of adult $A_{undiff}$ and $A_{diff}$ using FACS.**

(A) FACS strategy for spermatogonia isolation from adult mouse testes. E-Cadherin$^{+}$KIT$^{-}$CD9$^{high}$ and E-Cadherin$^{+}$KIT$^{+}$CD9$^{medium}$ subfractions were isolated as $A_{undiff}$ and $A_{diff}$ for downstream analyses. Average percentages of each gated fraction out of the total cells are shown. At least three independent experiments were analyzed for each genotype, and representative plots are shown.

(B) Immunostaining of cell markers in isolated spermatogonia. PLZF (green) is an undifferentiated spermatogonia marker, KIT (yellow) is a differentiating spermatogonia marker, and DDX4 (red) is a germ cell marker. Bars: 100 μm. At least three independent replicates were analyzed for each genotype, and representative images are shown.

(C) Quantifications of purities of isolated spermatogonia. Data are presented as mean values ± SD.

(D) Heatmap showing Pearson correlation values among each biological replicate in the WT spermatogonia RNA-seq dataset.

(E) Gene ontology analyses of the DEGs. Key GO terms of the DEGs are shown. P values were generated by Metascape using the two-sided hypergeometric test.

(F) Violin plots showing the expression level of previously defined marker genes for undifferentiated and differentiating spermatogonia (Green et al. 2018) in our WT $A_{undiff}$ and $A_{diff}$ bulk RNA-seq dataset. Two-tailed unpaired t-tests.

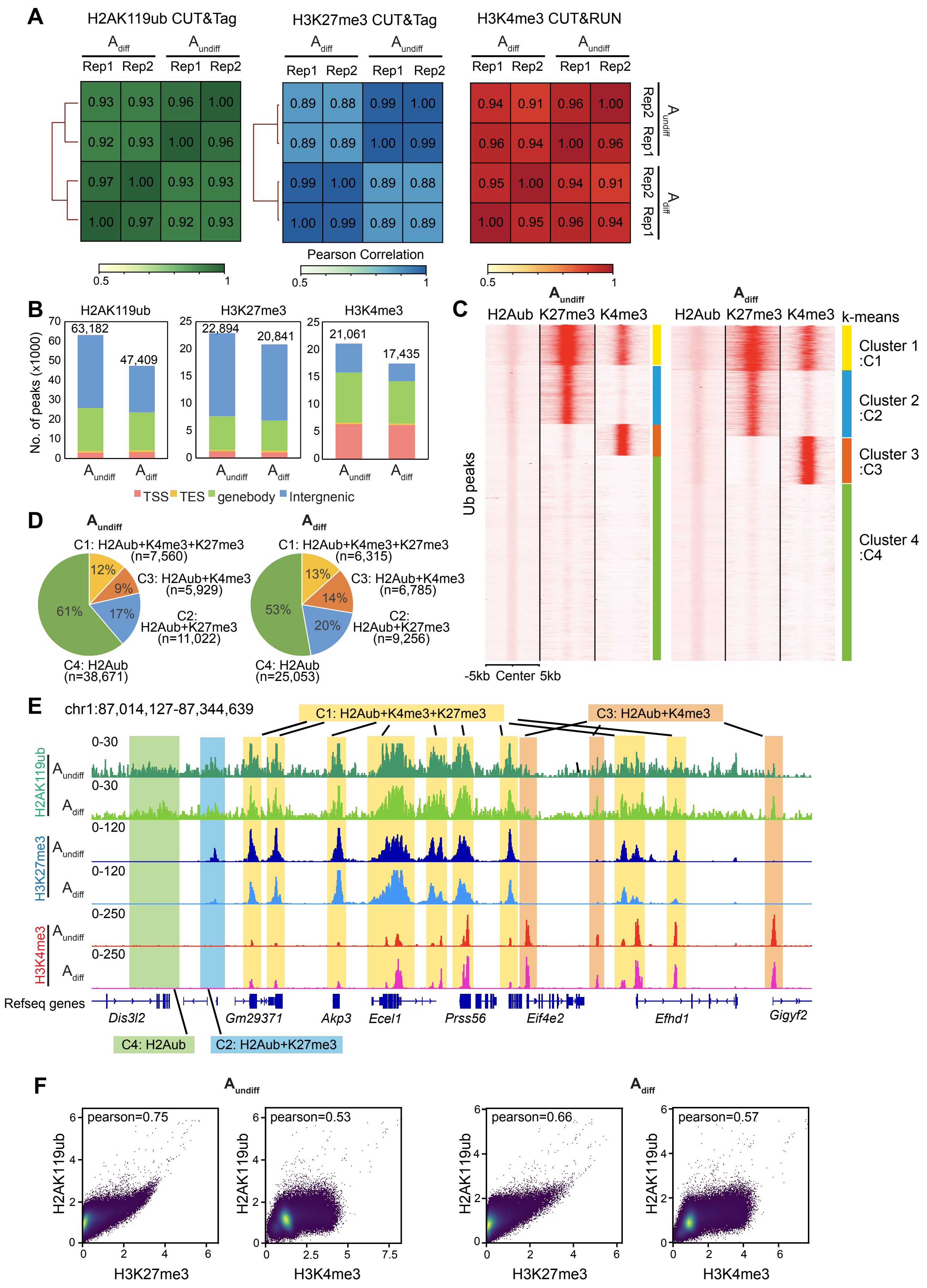

A
H2AK119ub CUT&Tag
H3K27me3 CUT&Tag
H3K4me3 CUT&RUN
A diff
A undiff
Rep1 Rep2 Rep1 Rep2
Pearson Correlation
0.5
1
B
H2AK119ub
H3K27me3
H3K4me3
No. of peaks (x1000)
63,182
47,409
22,894
20,841
21,061
17,435
A undiff
A diff
TSS TES genebody Intergnenic
C
A undiff
A diff
H2Aub K27me3 K4me3
k-means
Cluster 1 :C1
Cluster 2 :C2
Cluster 3 :C3
Cluster 4 :C4
Ub peaks
-5kb Center 5kb
D
C1: H2Aub+K4me3+K27me3 (n=7,560)
C3: H2Aub+K4me3 (n=5,929)
C2: H2Aub+K27me3 (n=11,022)
C4: H2Aub (n=38,671)
12%
9%
17%
61%
C1: H2Aub+K4me3+K27me3 (n=6,315)
C3: H2Aub+K4me3 (n=6,785)
C2: H2Aub+K27me3 (n=9,256)
C4: H2Aub (n=25,053)
13%
14%
20%
53%
E
chr1:87,014,127-87,344,639
C1: H2Aub+K4me3+K27me3
C3: H2Aub+K4me3
H2AK119ub
H3K27me3
H3K4me3
0-30
0-120
0-250
Refseq genes
Dis3l2
Gm29371
Akp3
Ecel1
Prss56
Eif4e2
Efhd1
Gigyf2
C4: H2Aub
C2: H2Aub+K27me3
F
A undiff
A diff
pearson=0.75
pearson=0.53
pearson=0.66
pearson=0.57
H2AK119ub
H3K27me3
H3K4me3

**Figure S2 Distributions of H2AK119ub, H3K27me3, and H3K4me3 in adult $A_{undiff}$ and $A_{diff}$.**
(A) Heatmaps showing the Pearson correlation values among each biological replicate in CUT&RUN/Tag data.
(B) Bar charts showing the number of H2AK119ub, H3K27me3, and H3K4me3 peaks and their genomic location in TSS, TES, gene body, and intergenic regions in $A_{undiff}$ and $A_{diff}$.
(C) Heatmaps showing H2AK119ub, H3K27me3, and H3K4me3 enrichment at all H2AK119ub peaks detected in each spermatogonial cell type with k-means clustering. Cluster 1 represents H2AK119ub (H2Aub)-covered regions that overlap both H3K27me3 (K27me3) and H3K4me3 (K4me3), Cluster 2 represents H2Aub-covered regions that overlap K27me3 only, Cluster 3 represents H2Aub-covered regions that overlap K4me3 only, Cluster 4 represents only H2Aub-covered regions.
(D) Pie charts showing the percentage of H2Aub-covered regions that overlap K27me3 or/and K4me3 detected in c in $A_{undiff}$ and $A_{diff}$.
(E) Representative track views showing different clusters of H2Aub-covered regions as indicated in C and D.
(F) Scatter plots showing the intensity (log (spike-in normalized reads count+1)) relationship between H2AK119ub and H3K27me3/H3K4me3 over 10-kb genomic bins in both $A_{undiff}$ and $A_{diff}$ . Pearson's correlation values are shown.

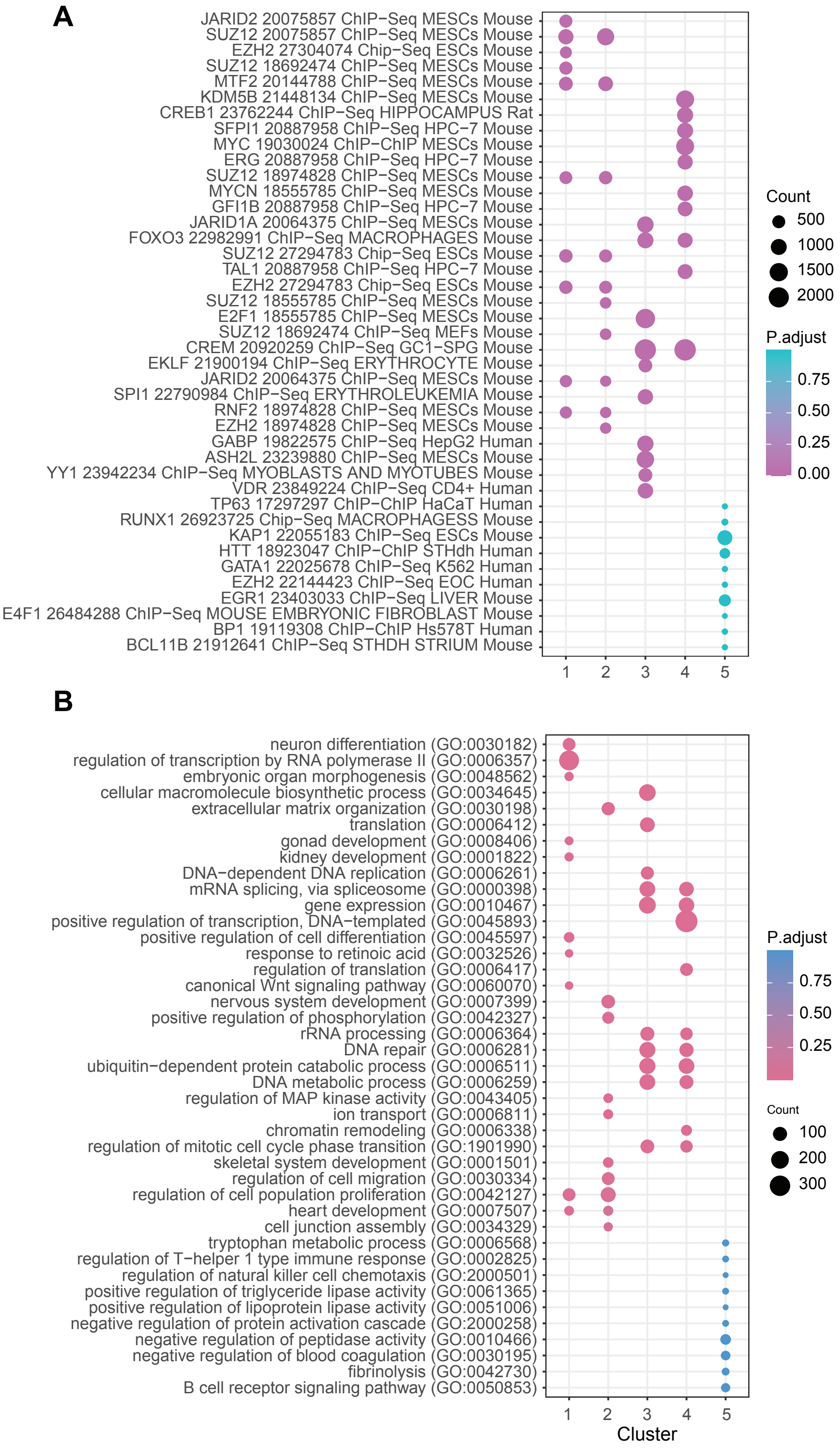
A
JARID2 20075857 ChIP−Seq MESCs Mouse
SUZ12 20075857 ChIP−Seq MESCs Mouse
EZH2 27304074 Chip−Seq ESCs Mouse
SUZ12 18692474 ChIP−Seq MESCs Mouse
MTF2 20144788 ChIP−Seq MESCs Mouse
KDM5B 21448134 ChIP−Seq MESCs Mouse
CREB1 23762244 ChIP−Seq HIPPOCAMPUS Rat
SFPI1 20887958 ChIP−Seq HPC−7 Mouse
MYC 19030024 ChIP−ChIP MESCs Mouse
ERG 20887958 ChIP−Seq HPC−7 Mouse
SUZ12 18974828 ChIP−Seq MESCs Mouse
MYCN 18555785 ChIP−Seq MESCs Mouse
GFI1B 20887958 ChIP−Seq HPC−7 Mouse
JARID1A 20064375 ChIP−Seq MESCs Mouse
FOXO3 22982991 ChIP−Seq MACROPHAGES Mouse
SUZ12 27294783 Chip−Seq ESCs Mouse
TAL1 20887958 ChIP−Seq HPC−7 Mouse
EZH2 27294783 Chip−Seq ESCs Mouse
SUZ12 18555785 ChIP−Seq MESCs Mouse
E2F1 18555785 ChIP−Seq MESCs Mouse
SUZ12 18692474 ChIP−Seq MEFs Mouse
CREM 20920259 ChIP−Seq GC1−SPG Mouse
EKLF 21900194 ChIP−Seq ERYTHROCYTE Mouse
JARID2 20064375 ChIP−Seq MESCs Mouse
SPI1 22790984 ChIP−Seq ERYTHROLEUKEMIA Mouse
RNF2 18974828 ChIP−Seq MESCs Mouse
EZH2 18974828 ChIP−Seq MESCs Mouse
GABP 19822575 ChIP−Seq HepG2 Human
ASH2L 23239880 ChIP−Seq MESCs Mouse
YY1 23942234 ChIP−Seq MYOBLASTS AND MYOTUBES Mouse
VDR 23849224 ChIP−Seq CD4+ Human
TP63 17297297 ChIP−ChIP HaCaT Human
RUNX1 26923725 Chip−Seq MACROPHAGESS Mouse
KAP1 22055183 ChIP−Seq ESCs Mouse
HTT 18923047 ChIP−ChIP STHdh Human
GATA1 22025678 ChIP−Seq K562 Human
EZH2 22144423 ChIP−Seq EOC Human
EGR1 23403033 ChIP−Seq LIVER Mouse
E4F1 26484288 ChIP−Seq MOUSE EMBRYONIC FIBROBLAST Mouse
BP1 19119308 ChIP−ChIP Hs578T Human
BCL11B 21912641 ChIP−Seq STHDH STRIUM Mouse
1
2
3
4
5
Count
500
1000
1500
2000
P.adjust
0.75
0.50
0.25
0.00
B
neuron differentiation (GO:0030182)
regulation of transcription by RNA polymerase II (GO:0006357)
embryonic organ morphogenesis (GO:0048562)
cellular macromolecule biosynthetic process (GO:0034645)
extracellular matrix organization (GO:0030198)
translation (GO:0006412)
gonad development (GO:0008406)
kidney development (GO:0001822)
DNA−dependent DNA replication (GO:0006261)
mRNA splicing, via spliceosome (GO:0000398)
gene expression (GO:0010467)
positive regulation of transcription, DNA−templated (GO:0045893)
positive regulation of cell differentiation (GO:0045597)
response to retinoic acid (GO:0032526)
regulation of translation (GO:0006417)
canonical Wnt signaling pathway (GO:0060070)
nervous system development (GO:0007399)
positive regulation of phosphorylation (GO:0042327)
rRNA processing (GO:0006364)
DNA repair (GO:0006281)
ubiquitin−dependent protein catabolic process (GO:0006511)
DNA metabolic process (GO:0006259)
regulation of MAP kinase activity (GO:0043405)
ion transport (GO:0006811)
chromatin remodeling (GO:0006338)
regulation of mitotic cell cycle phase transition (GO:1901990)
skeletal system development (GO:0001501)
regulation of cell migration (GO:0030334)
regulation of cell population proliferation (GO:0042127)
heart development (GO:0007507)
cell junction assembly (GO:0034329)
tryptophan metabolic process (GO:0006568)
regulation of T−helper 1 type immune response (GO:0002825)
regulation of natural killer cell chemotaxis (GO:2000501)
positive regulation of triglyceride lipase activity (GO:0061365)
positive regulation of lipoprotein lipase activity (GO:0051006)
negative regulation of protein activation cascade (GO:2000258)
negative regulation of peptidase activity (GO:0010466)
negative regulation of blood coagulation (GO:0030195)
fibrinolysis (GO:0042730)
B cell receptor signaling pathway (GO:0050853)
1
2
3
4
5
Cluster
P.adjust
0.75
0.50
0.25
Count
100
200
300

**Figure S3  H2AK119ub, H3K27me3, and H3K4me3 target genes in adult spermatogonia.**

(A and B)  Dot plots of ChIP-x Enrichment Analysis (ChEA) (A) and Gene Ontology Biological Process (GO-BP) term enrichment (B). The Top 10 enriched GO-BP terms and predicted transcription factors in the gene set of each k-means cluster (identified in Figure 2F). ChEA enrichment shows the transcription factors and the cell and animal types used in the profiling experiments. Colors indicate the adjusted p-value using the Benjamini-Hochberg method for correction for multiple hypotheses testing, and dots size is proportional to gene count, the number of genes enriched in the gene set library.

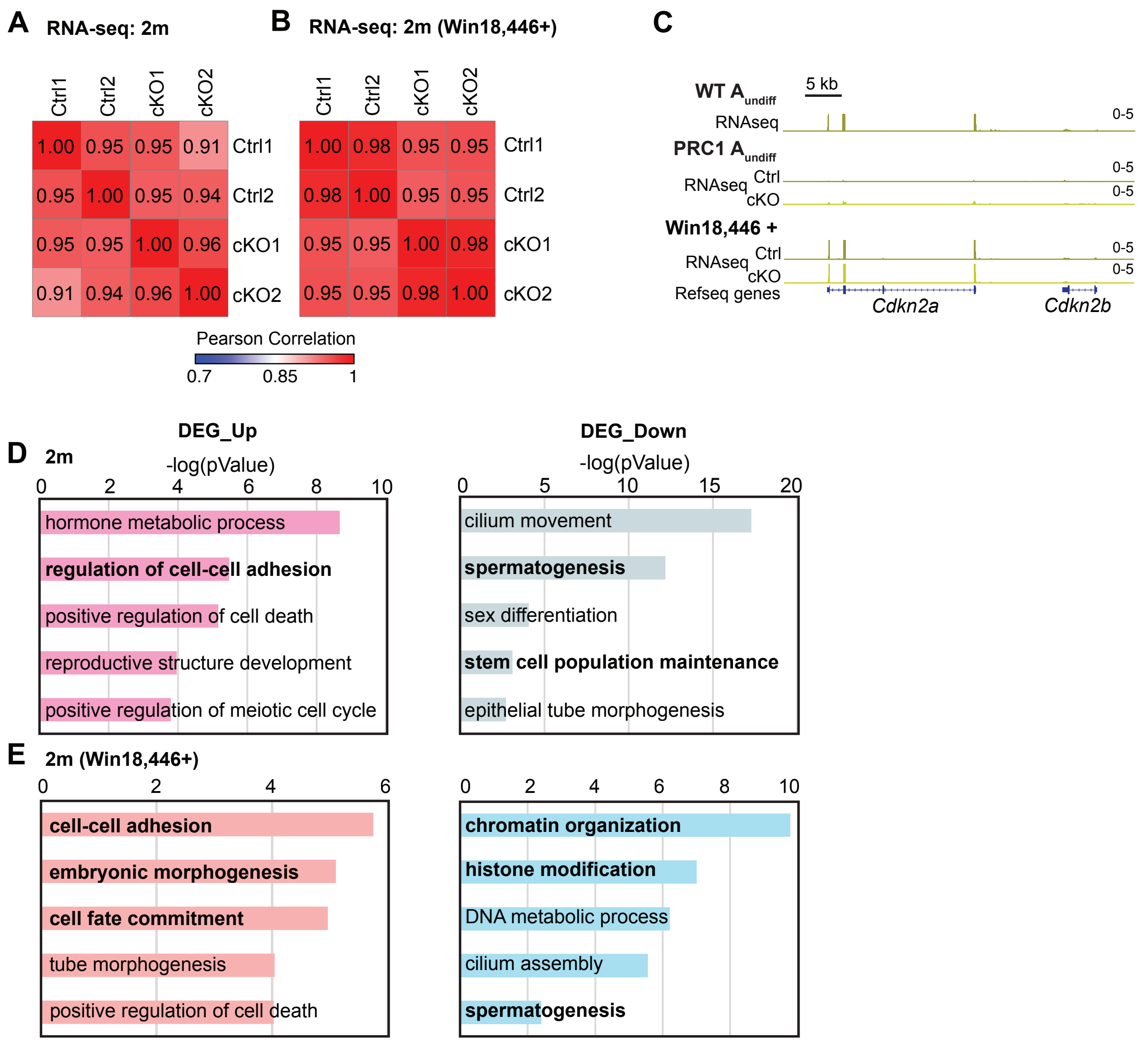


**Figure S4 Gene expression in PRC1ctrl and PRC1cKO $A_{undiff}$.**
(A and B) Heatmaps showing the Pearson correlation values among biological replicates in PRC1ctrl and PRC1cKO $A_{undiff}$ RNA-seq dataset.
(C) Track views of the *Cdkn2a/b* loci showing the RNA-seq read density in $A_{undiff}$ of indicated genotypes. TPM ranges are shown in the upper right.
(D and E) Gene ontology analyses of the DEGs. Key GO terms of the DEGs are shown. P values were generated by Metascape using the two-sided hypergeometric test.

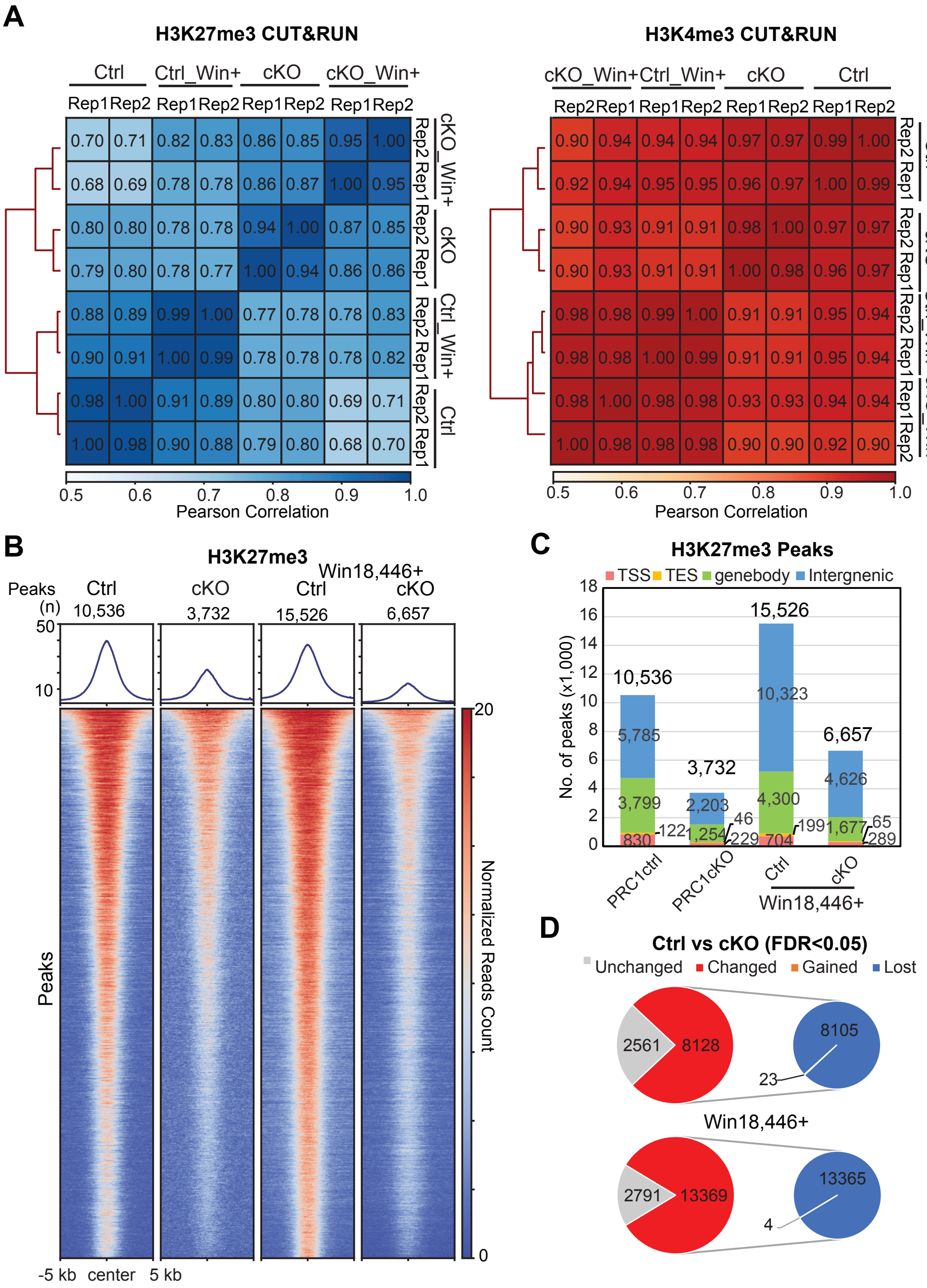


**Figure S5 PRC1-dependent regulation of PRC2-mediated H3K27me3 in $A_{undiff}$.**

(A) Heatmaps with hierarchical clustering showing the Pearson correlation values among each biological replicate in PRC1ctrl and PRC1cKO $A_{undiff}$ CUT&RUN data.

(B) Heatmaps and average tag density plots showing H3K27me3 peaks in PRC1ctrl and PRC1cKO $A_{undiff}$ at 2 mo with or without Win18,446 treatment.

(C) Bar charts showing H3K27me3 peak number and genomic distribution in $A_{undiff}$ of indicated genotypes.

(D) Pie charts showing H3K27me3 peaks change upon PRC1 inactivation in $A_{undiff}$ at 2 mo with or without Win18,446 treatment.

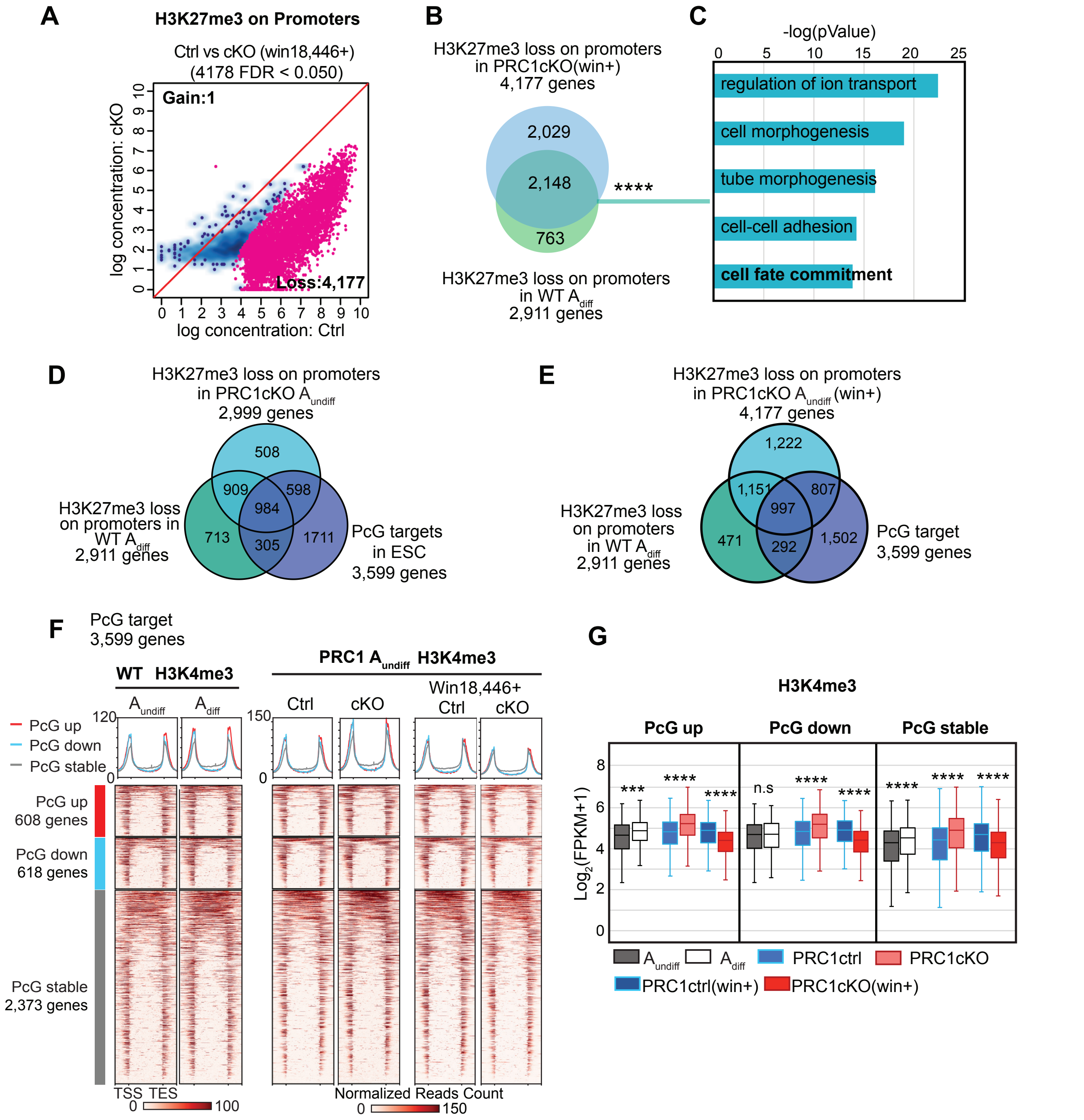


**Figure S6 Polycomb shields the lineage-specific developmental program in adult spermatogonia.**

(A) Scatter plot comparing H3K27me3 enrichment at promoters (TSS ± 2 kb) in PRC1ctrl and PRC1cKO $A_{undiff}$ at 2 mo after win18,446 treatment. Red dots represent genes with differential H3K27me3 enrichment at promoters (FDR < 0.05).

(B) Overlap between H3K27me3-lost promoters in PRC1cKO $A_{undiff}$ after win18,446 treatment and H3K27me3-lost promoters in WT $A_{diff}$. ****$P = 5.85 \times 10^{-1165}$, two-sided hypergeometric test.

(C) Key GO enrichments in the overlap detected in B.

(D and E) Venn diagrams showing overlaps among H3K27me3-lost promoters in PRC1cKO $A_{undiff}$ with or without Win18,446 treatment, H3K27me3-lost promoters in WT $A_{diff}$ and classic Polycomb targets in ESCs.

(F) Heatmaps and average tag density plots of H3K4me3 enrichment at genic regions (TSS-TES ± 2 kb) on different Polycomb-targets groups in WT $A_{undiff}$ and $A_{diff}$ or PRC1 $A_{undiff}$ of indicated genotypes. The bars below the heatmaps represent signal intensity, and the numbers represent spike-in normalized read counts.

(G) Box-and-whisker plots showing H3K4me3 enrichment at promoters (TSS ± 2 kb, log2-transformed FPKM) in the corresponding gene groups shown in F. Central bars represent medians, the boxes encompass 50% of the data points, and the whiskers indicate 90% of the data points. n.s, not significant, *** $P < 0.001$, **** $P < 2.2 \times 10^{-6}$; two-tailed unpaired t-tests.

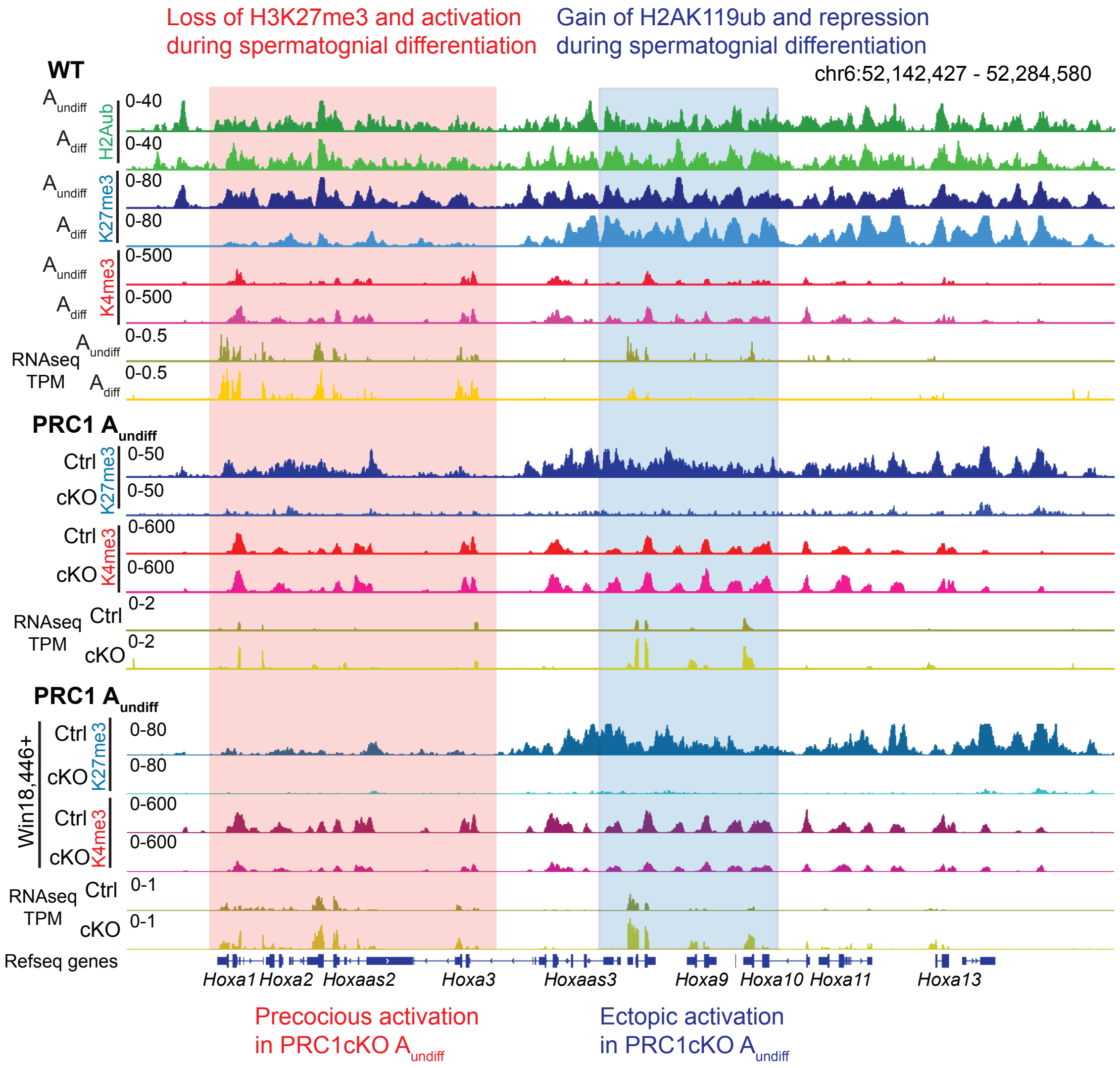


**Figure S7 Track views of the *Hoxa* loci.**
Track views of the *Hoxa* loci showing H2AK119ub, H3K27me3, and H3K4me3 enrichment and RNA-seq reads in WT $A_{undiff}$ and $A_{diff}$ or PRC1 $A_{undiff}$ of indicated genotypes. Data ranges are shown in the upper left.

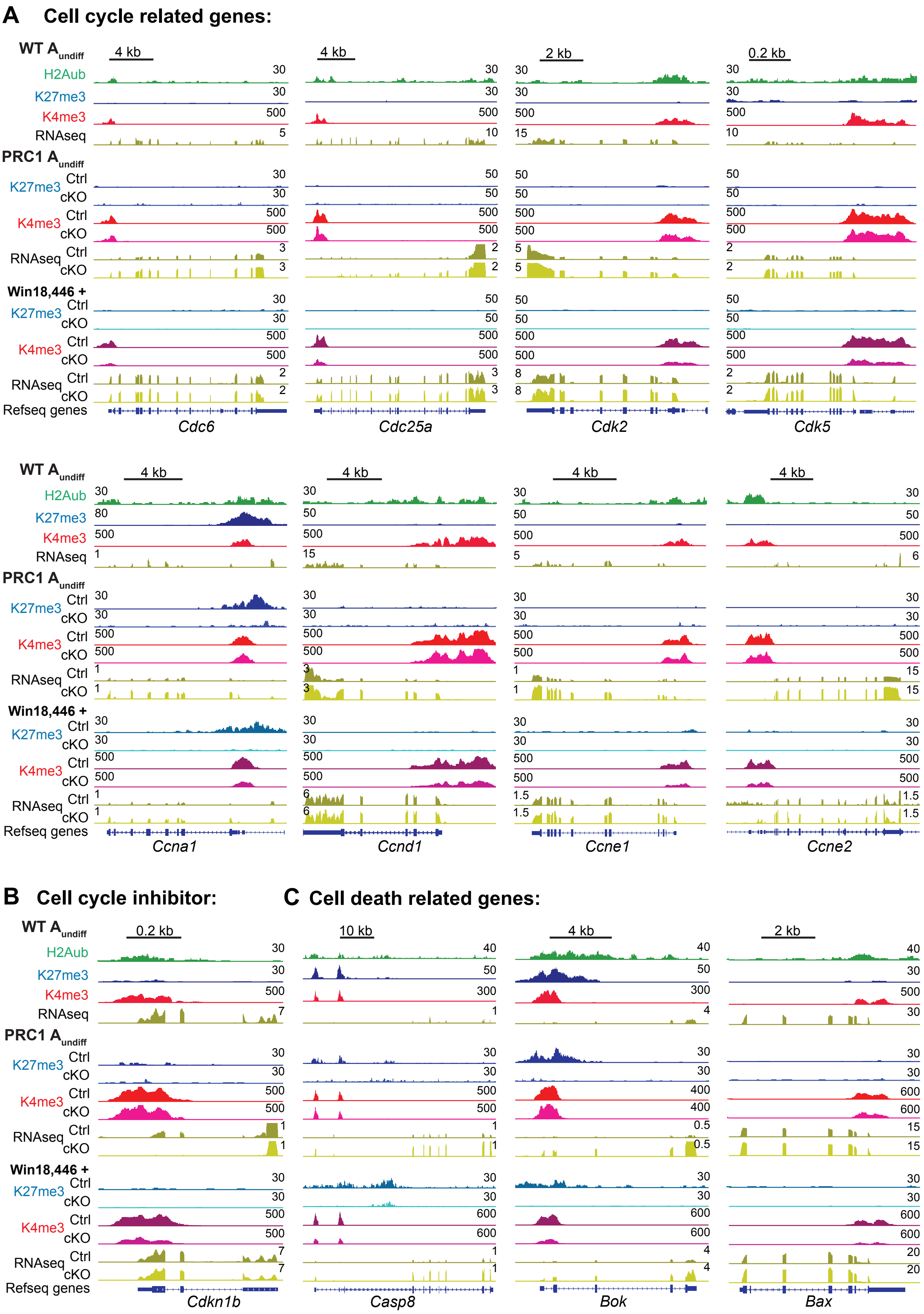

A Cell cycle related genes:
WT A undiff
H2Aub
K27me3
K4me3
RNAseq
PRC1 A undiff
K27me3 Ctrl cKO
K4me3 Ctrl cKO
RNAseq Ctrl cKO
Win18,446 +
K27me3 Ctrl cKO
K4me3 Ctrl cKO
RNAseq Ctrl cKO
Refseq genes
4 kb
2 kb
0.2 kb
Cdc6
Cdc25a
Cdk2
Cdk5
Ccna1
Ccnd1
Ccne1
Ccne2
B Cell cycle inhibitor:
Cdkn1b
C Cell death related genes:
10 kb
Casp8
Bok
Bax

**Figure S8 PRC1 deficiency activates cell cycle- and cell death-related genes.**
(A, B, and C) Track views of cell cycle-related gene loci (A, B) and cell death-related gene loci (C) showing H2AK119ub, H3K27me3, and H3K4me3 enrichment and RNA-seq reads in $A_{undiff}$ of indicated genotypes. Data ranges (maximum numbers only) are shown in tracks.

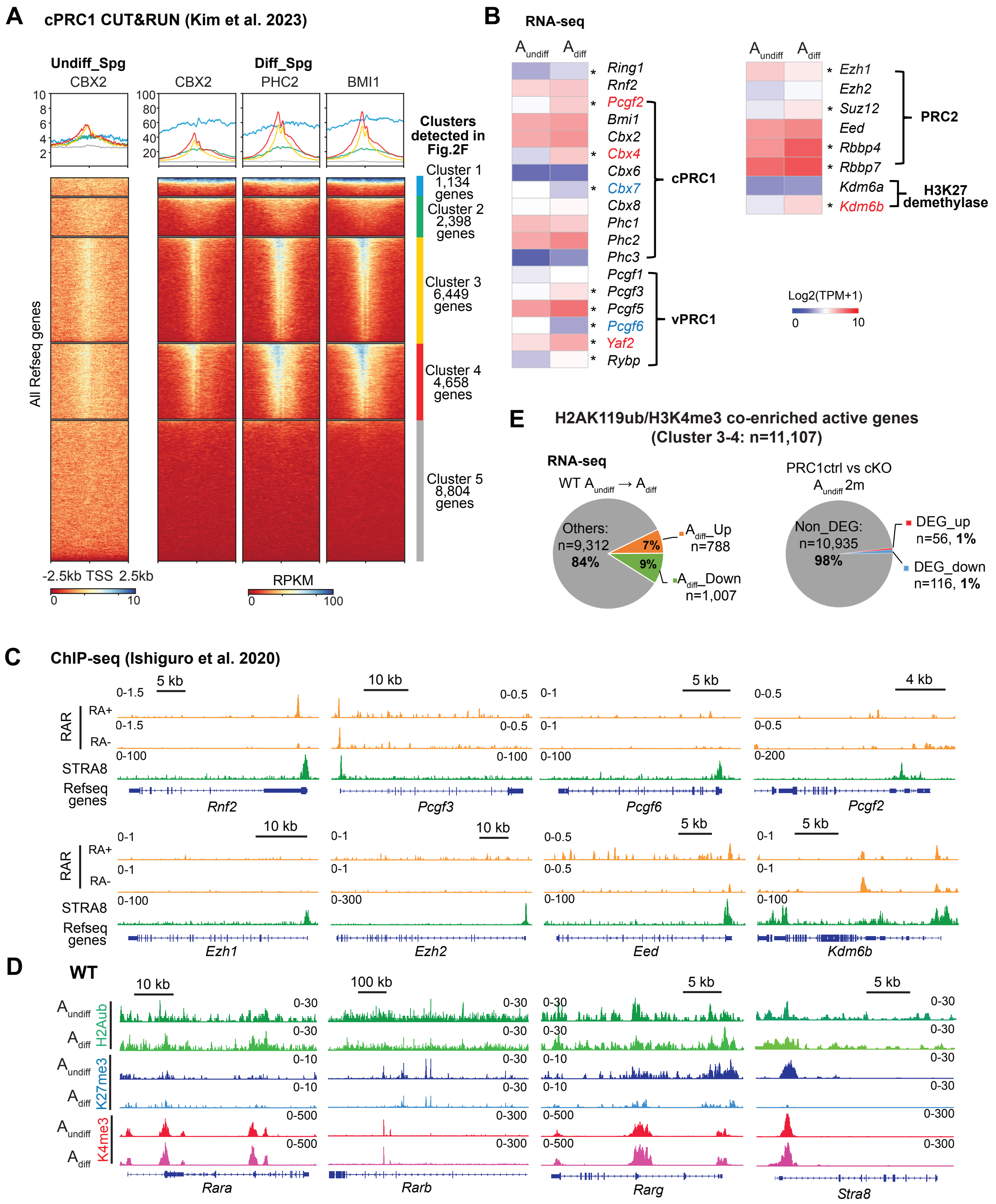

A
cPRC1 CUT&RUN (Kim et al. 2023)
Undiff_Spg
CBX2
Diff_Spg
CBX2
PHC2
BMI1
Clusters detected in Fig.2F
Cluster 1 1,134 genes
Cluster 2 2,398 genes
Cluster 3 6,449 genes
Cluster 4 4,658 genes
Cluster 5 8,804 genes
All Refseq genes
-2.5kb TSS 2.5kb
RPKM
B
RNA-seq
Ring1
Rnf2
Pcgf2
Bmi1
Cbx2
Cbx4
Cbx6
Cbx7
Cbx8
Phc1
Phc2
Phc3
cPRC1
Pcgf1
Pcgf3
Pcgf5
Pcgf6
Yaf2
Rybp
vPRC1
Ezh1
Ezh2
Suz12
Eed
Rbbp4
Rbbp7
PRC2
Kdm6a
Kdm6b
H3K27 demethylase
Log2(TPM+1)
E
H2AK119ub/H3K4me3 co-enriched active genes (Cluster 3-4: n=11,107)
RNA-seq
Others: n=9,312 84%
7%
9%
n=788
n=1,007
PRC1ctrl vs cKO
Non_DEG: n=10,935 98%
DEG_up n=56, 1%
DEG_down n=116, 1%
C
ChIP-seq (Ishiguro et al. 2020)
RAR
RA+
RA-
STRA8
Refseq genes
Rnf2
Pcgf3
Pcgf6
Pcgf2
Ezh1
Ezh2
Eed
Kdm6b
D
WT
H2Aub
K27me3
K4me3
Rara
Rarb
Rarg
Stra8

**Figure S9 Data analysis related to Discussion.**
(A) Heatmaps and average tag density plots of k-means clusters (defined in Fig.2F) showing components of cPRC1:CBX2, PHC2, and BMI1 enrichment (Kim et al. 2023) at promoter regions (TSS ± 2.5 kb) in WT undifferentiated spermatogonia (Undiff_Spg: THY1$^{+}$ & ITGa6$^{+}$) and differentiating spermatogonia (Diff_Spg: KIT$^{+}$). The color keys represent signal intensity and the numbers represent RPKM values.
(B) Heatmap showing gene expression of representative Polycomb components and H3K27 demethylases in WT Aundiff and Adiff populations. *, FDR< 0.05. Gene names are color-coded to indicate DEG status (Log2FoldChange >1, FDR< 0.05); Blue: downregulated DEGs; Red: upregulated DEGs.
(C) Track views of gene loci representing Polycomb components and H3K27 demethylase Kdm6b gene loci showing RA receptor (pan-RAR) occupancy in GS cells with or without RA treatment and STRA8 binding in testes (Ishiguro et al. 2020). RAR ChIP-seq data ranges represent fold enrichment calculated using local lambda values (downloaded from GSE116798). STRA8 ChIP-seq data ranges represent RPKM values.
(D) Track views of RARs and *Stra8* loci showing H2AK119ub, H3K27me3, and H3K4me3 enrichment in WT Aundiff and Adiff. Data ranges are shown in the tracks. Notably, RAR-STRA8 and Polycomb can be direct targets of each other, suggesting the Polycomb system and RA-STRA8 signaling may form a regulatory loop that coordinates functions of each other.
(E) Pie charts showing DEG distributions in H2AK119ub/H3K4me3 co-enriched genes (Cluster 3-4, identified in Fig. 2F) in WT spermatogonia and PRC1 Aundiff RNA-seq, respectively. The percentages and number (n) of up-and down-regulated DEGs and non-DEGs are indicated.
